# A mechanical G2 checkpoint controls epithelial cell division through E-cadherin-mediated regulation of Wee1-Cdk1

**DOI:** 10.1101/2021.11.29.470352

**Authors:** Lisa Donker, Marjolein J. Vliem, Helena Canever, Manuel Gómez-González, Miquel Bosch-Padrós, Willem-Jan Pannekoek, Xavier Trepat, Nicolas Borghi, Martijn Gloerich

**Author notes:** Correspondence to Martijn Gloerich.

## Abstract

Epithelial cell divisions must be tightly coordinated with cell loss to preserve epithelial integrity. However, it is not well understood how the rate of epithelial cell division adapts to changes in cell number, for instance during homeostatic turnover or upon wounding of epithelia. Here, we show epithelial cells sense local cell density through mechanosensitive E-cadherin adhesions to control G2/M cell cycle progression. We demonstrate that tensile forces on E-cadherin adhesions are reduced as local cell density increases, which prompts the accumulation of the G2 checkpoint kinase Wee1. This elevated abundance of Wee1 results in inhibitory phoshorylation of Cdk1, and thereby establishes a pool of cells that is temporarily halted in G2-phase. Importantly, these cells are readily triggered to divide upon epithelial wounding, due to the consequent increase in intercellular forces and resulting degradation of Wee1. Our data thus demonstrate that epithelial cell division is controlled by a mechanical G2 checkpoint, which is regulated by cell density-dependent intercellular forces sensed and transduced by E-cadherin adhesions.

## Introduction

Epithelia form essential, protective barriers covering the body surface and internal organs, yet display high rates of cellular turnover. To preserve the integrity of the epithelial barrier while at the same time preventing tissue overgrowth, epithelial cell division must be tightly coordinated with cell loss. This requires epithelia to respond to fluctuations in cell number, for instance during homeostatic turnover or due to wounding, and adjust their rate of cell division accordingly. Disruption of this proliferative response underlies epithelial barrier disorders (Eming et al., 2014), and epithelial cancers in which density-dependent inhibition of proliferation is typically lost (McClatchey and Yap, 2012). Understanding how epithelial homeostasis is achieved and maintained requires identification of the mechanisms by which epithelia sense changes in cell number and transmit this information to adapt division rate.

Epithelial cell division can be induced by activation of quiescent cells that have exited the cell cycle, and it has previously been demonstrated that cells residing in G0/G1 are triggered to re-enter the cell cycle following wounding of the epithelial layer (Bornes et al., 2021; Streichan et al., 2014). In most epithelial tissues, cells with proliferative capacity are already actively cycling during homeostasis to drive tissue self-renewal (Li and Clevers, 2010). The division rate of these cells is presumed to be independent of external regulation, as it has been a prevailing dogma that following G1/S transition, progression through the cell cycle becomes irresponsive to extrinsic cues (Pardee, 1989). However, recent findings in invertebrate germline and neural stem cells revealed that progression through later cell cycle stages can be modulated by nutrient signaling (McKeown and Cline, 2019; Otsuki and Brand, 2018; Seidel and Kimble, 2015). Yet, how extrinsic signals influence progression through later stages of the cell cycle and thereby the rate of cell division in epithelia is not well understood.

Extrinsic regulation of cell proliferation not only involves biochemical signals (e.g. nutrients and growth factors), but also mechanical forces originating from neighboring cells and the extracellular environment (Godard and Heisenberg, 2019; LeGoff and Lecuit, 2016). Cells possess a repertoire of mechanosensor proteins that can transduce these mechanical cues into an intracellular response, involving force-dependent changes in their conformation and subsequent induction of downstream signaling pathways (Swaminathan and Gloerich, 2021). As such, mechanotransduction by cadherin- and integrin-based adhesions that anchor cells to each other and the surrounding tissue, respectively, contribute to the regulation of cell cycle exit and re-entry (Aragona et al., 2013; Benham-Pyle et al., 2015; Dutta et al., 2018; Ibar et al., 2018; Rauskolb et al., 2014). Also the force-sensitive calcium channel Piezo1 is linked to epithelial cell proliferation and was recently shown to play a role in mitotic entry following the ectopic application of mechanical stretch (Gudipaty et al., 2017). Nonetheless, the forces that cells experience during epithelial homeostasis, and how these forces regulate cell cycle progression, remain poorly understood.

Here, we show that epithelial cells sense changes in cell number through coinciding changes in intercellular forces, which are transduced by E-cadherin adhesions to control a cell density-dependent G2 checkpoint. This checkpoint establishes a pool of cells that is halted in the G2-phase of the cell cycle in dense epithelia, mediated by the force-sensitive regulation of the Wee1 kinase that maintains the mitotic gatekeeper Cdk1 in an inactive state. Increased tension on E-cadherin following elevation of intercellular forces, for instance during epithelial expansion upon wounding, triggers rapid degradation of Wee1 and subsequent mitotic entry of these G2-halted cells. Our data thus demonstrate that epithelia coordinate cell divisions with local cell density through a mechanical G2 checkpoint regulated by E-cadherin mechanotransduction.

## Results

### Rapid induction of mitotic entry during epithelial expansion following wounding

To investigate how epithelial cell divisions are coordinated with cell loss, we analyzed cell division rate during epithelial expansion upon wounding of monolayers of Madin-Darby Canine Kidney (MDCK) cells that recapitulate the organization and function of a single-layered epithelial tissue. Characterization of MDCK monolayers 24h after plating in wells within silicone (polydimethylsiloxane; PDMS) stencils demonstrated they formed confluent monolayers starting at ~1000 cells / mm^2^, and remained almost fully Ki67-positive and thus actively cycling up to a density of ~4500 cells / mm^2^ (**Figs. 1a, 1b**). We therefore defined monolayers of 2600-4500 cells / mm^2^ as high density (yet still actively cycling) monolayers, 1000 – 2200 cells / mm^2^ as low density monolayers, and >4500 cells / mm^2^ as monolayers undergoing contact inhibition of proliferation and exiting the cell cycle. Next, we mimicked the increase in available surface that occurs upon epithelial wounding by removal of the PDMS stencil, which allows cells to migrate freely into the newly available space (Poujade et al., 2007). This approach recapitulates the changes in cell motility and cell density, and concurrent epithelial expansion that occur upon epithelial wounding, in the absence of cell damage induced by traditional wounding assays (Poujade et al., 2007). Strikingly, in monolayers cultured at a high density, we observed a transient burst of mitotic events within 1-2 hours after stencil removal (**Figs. 1d, 1e**). The number of mitotic cells returned to basal levels within 4 hours after stencil removal (**Fig. 1e**). In low density monolayers, the number of mitotic events prior to stencil removal was higher than in high density monolayers, but remained unaltered during epithelial expansion (**Figs. 1d, 1e**). Expectedly, also contact inhibited monolayers (> 4500 cells / mm^2^) did not show an elevation of mitotic events within the first hours following stencil removal (**Fig. S1a**). These data indicate that dense monolayers of actively cycling cells rapidly induce a pool of cells to enter mitosis and divide during epithelial expansion following wounding.

**Figure 1.**
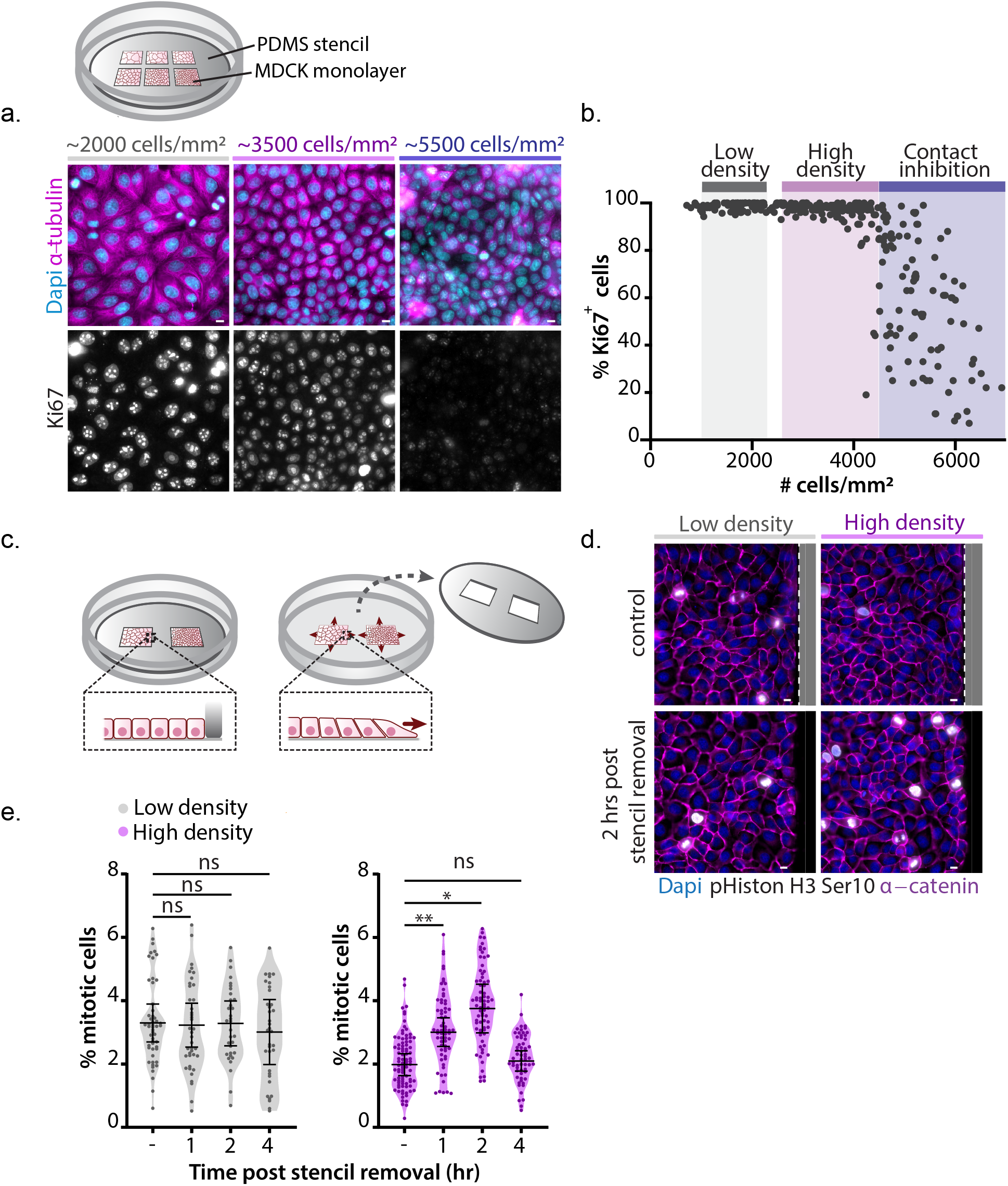
Rapid induction of mitotic entry during epithelial expansion following wounding. **(a)** Top: schematic image of wells within a (polydimethylsiloxane) PDMS stencil that is placed on a Collagen I-coated glass surface, with different densities of MDCK cells plated in each well. Within these wells all formed monolayers are exposed to the same culture medium and thus identical concentrations of nutrients and growth factors. Bottom: Immunostaining of MDCK monolayers at various cell densities for the proliferation marker Ki67 together with α-tubulin and Dapi. **(b)** Quantification of the percentage of Ki67^+^ cells in MDCK monolayers grown at various cell densities. Most cells remain Ki67-positive until monolayers reach a density of ~4500 cells/mm^2^, which indicates the start of contact inhibition of proliferation. The range of densities considered low density monolayers (1000 - 2200 cells/mm^2^), high density monolayers (2600 – 4500 cells/mm^2^), and contact-inhibited density (CIP) (>4500 cells/mm^2^) are indicated in grey, magenta and blue, respectively (and used from here onwards in the manuscript). Data were pooled from 3 independent experiments, with > 50 monolayer regions per experiment. Scale bars represent 10 μm. **(c)** Schematic image of the stencil removal assay to mimic epithelial expansion following wounding, in which removal of a PDMS stencil allows cells to migrate freely onto the newly available surface. **(d)** Immunostaining of MDCK monolayers (grown at low and high density (see Fig. 1b) without and 2 hours after induction of epithelial expansion by removal of the PDMS stencil, for phospho-Histon H3 Ser10 together with α-catenin and Dapi. The PDMS stencil is indicated in grey, and the dotted line indicates the border between the stencil and the monolayers of cells. Scale bars represent 10 μm. **(e)** Quantification of the percentage of mitotic cells in unperturbed MDCK monolayers grown at low (grey) and high (magenta) monolayer density before and 1, 2 and 4 hrs post stencil removal. n > 30 monolayer regions per condition. Data were pooled from 4 independent experiments. Black bars represent the mean and SD of the individual experiments. *P = 0.023; **P = 0.007, ns = not significant; paired t-test.

### G2-phase of the cell cycle is prolonged in dense epithelial monolayers

We next investigated why induction of mitotic events during wounding-induced epithelial expansion specifically occurred in high density epithelia. As this mitotic response occurred within 1-2 hours, this suggests that the cells triggered to enter mitosis were in the G2-phase of the cell cycle. To test this, we generated MDCK cells stably expressing the FUCCI4 reporter (Bajar et al., 2016), which allows for visualization of all four cell cycle phases (**Fig. 2a**). This showed that cells entering mitosis during expansion of dense MDCK-FUCCI4 monolayers indeed resided in G2-phase prior to stencil release, as indicated by the presence of Clover-Geminin(1-110) and concurrent absence of mTurquoise2-SLBP(18-126) expression (**Fig. 2b**).

**Figure 2.**
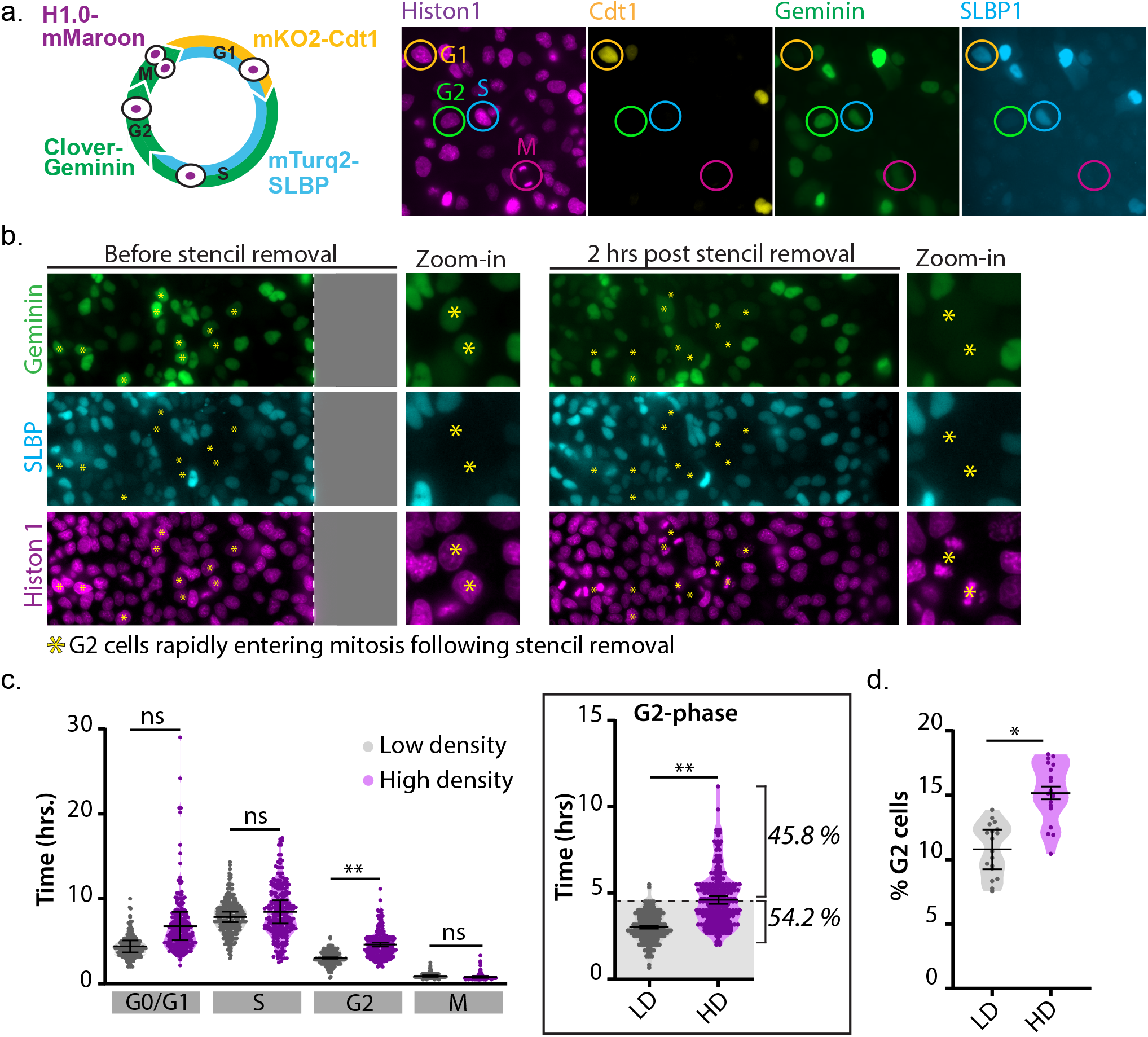
Prolongation of G2-phase in dense epithelial monolayers. **(a)** Schematic image of the FUCCI4 reporter (Bajar et al., 2016), showing differential expression of mKO-Cdt1(30-120), mTurquoise2-SLBP(18-126), Clover-Geminin(1-110) throughout the cell cycle, together with H1.0-mMaroon to visualize chromosome condensation in mitosis. **(b)** Visualization of cell cycle stages using the FUCCI4 reporter in dense MDCK monolayers before and 2 hours after induction of epithelial expansion by removal of the (polydimethylsiloxane) PDMS stencil. All cells that enter mitosis during epithelial expansion following stencil removal (asterisk) were in the G2-phase of the cell cycle prior to stencil removal (indicated by expression of Clover-Geminin(1-110) and absence of mTurquoise2-SLBP(18-126)). Scale bars represent 10 μm. **(c)** Left: Quantification of the duration of the different cell cycle phases (hrs) of cells grown at low (grey) and high (magenta) monolayer density. Note that G0- and G1-phase cannot be distinguished by the FUCCI4 reporter, although at the cell densities used in this experiment very few cells will be in G0 (Figs. 1a, 1b). While the average length of G1-phase does not significantly change, there is a pool of cells (22.2 ± 16.07 %) at high density with a prolonged (>8 hrs) G1-phase (Fig. S1c). Right: inset showing the duration of G2-phase (hrs) in monolayers grown at low (grey) and high (magenta) density, in which the 99 percentile of the duration of G2 in low density monolayers is indicated (4.5 hrs, dashed line), with 45.6 ± 6.2% of cells in high density monolayers showing a prolonged G2 duration longer than 4.5 hrs. n = 225 cells per condition. Data were pooled from 3 independent experiments. Black bars represent the mean and SD of the individual experiments. **P < 0.0034; ns = not significant; paired t-test. **(d)** Quantification of the percentage of G2-phase cells in low density and high density monolayers. n = 18 monolayer regions per condition. Data were pooled from 3 independent experiments. Black bars represent the mean and SD of the individual experiments. *P = 0.0275; paired t-test.

Because the mitotic response during expansion of MDCK monolayers occurred only in dense monolayers (**Figs. 1d, 1e**), we hypothesized that high density monolayers contain a larger population of G2-cells that is able to respond to wounding compared to low density monolayers. To investigate this, we determined the length of G2 and other phases of the cell cycle by live-imaging MDCK-FUCCI4 monolayers over time. In low density monolayers, the average duration of G2-phase was 3.0 ± 0.8 hours, with 99% of the cells completing G2-phase within 4.5 hours (**Fig. 2c**). At high cell density the duration of G2-phase was significantly prolonged (**Fig. 2c**), which strongly anti-correlated with the local distance between cell nuclei (**Fig. S1b**). While the average increase in G2 length at high density was ~1.6 hours (from 3.0 ± 0.8 to 4.6 ± 1.5 hrs), a significant population of cells (45.8 ± 6.2 %) displayed a prolonged G2 duration, ranging from 4.5 up to ~10 hrs (**Fig. 2c**).

The average duration of the G0/G1 phases (which cannot be separated from each other by the FUCCI4 reporter) and S-phase did not significantly change between both monolayer densities (**Fig. 2c**). However, high density monolayers did contain a substantial number of cells (22.2 ± 16.07 %) with a prolonged G0/G1 length compared to low density monolayers (**Figs. 2c, S1c**). We did not observe a clear correlation between the length of G0/G1- and G2-phases within individual cells (**Fig. S1d**), and the majority of cells with a prolonged (> 4.5 hours) G2-phase (65.9 ± 20.4 %) did not show a similar G0/G1 prolongation earlier in their cell cycle (**Fig. S1e**). This suggests that, although both G0/G1 and G2 length are increased at high cell density, the underlying mechanisms by which cell density influences these phases of the cell cycle are likely distinct.

As a result of the G2 prolongation, dense epithelia contained a significantly increased number of cells in G2 (average of 15.2%) compared to low density monolayers (average of 10.8%) (**Fig. 2d**). Importantly, to exclude that the extended G2 length and concurrent increase in the pool of G2 cells at high density is a consequence of increased DNA damage, we assessed the amount of DNA double-strand breaks by immunostaining for yH2AX (**Fig. S1f**). This showed the absence of yH2AX foci at high density, which in contrast were clearly present following UV-irradiation, indicating that the density-dependent G2 regulation occurs independently of a DNA damage response.

Taken together, our data indicate that in epithelial layers the duration of G2-phase is influenced by local cell density. This results in a larger pool of cells in G2-phase in dense epithelia, which can be triggered to enter mitosis during epithelial expansion following wounding.

### Density-dependent G2/M regulation requires mechanotransduction through E-cadherin adhesions

Having established that G2-phase is prolonged at high epithelial density, we next investigated how cells sense differences in monolayer density to adapt G2 length. As the density of epithelial monolayers increases, the motility of epithelial cells gradually declines, as visualized by particle image velocimetry (PIV) analysis (**Figs. 3a, 3b**; ref. (Petitjean et al., 2010)). Importantly, we found that this gradual decrease in motility occurred across densities at which we observed prolongation of G2-phase, and plateaued at a density of ~4500 cells / mm^2^ (**Fig. 3b**). These reduced intercellular movements at high monolayer density could potentially impact the level of forces that cells exert on each other. To map these intercellular forces we performed monolayer stress microscopy (Tambe et al., 2011) in monolayers at various densities, showing a strong reduction in intercellular tensions as cells reach high density (**Figs. 3c, 3d**), in line with the density-dependent reduction of cell motility (**Figs. 3a, 3b**). Intercellular forces are transmitted by E-cadherin adhesions, which form homotypic interactions between neighboring cells and are linked to the actin cytoskeleton through associated catenin proteins (Leckband and de Rooij, 2014). To test whether an increase in cell density results in a reduction in tension on the E-cadherin complex, we made use of an E-cadherin tension sensor (E-cadherin TsMod), in which tensile forces separate the mTFP1/YFP FRET pair placed in the cytosolic tail of E-cadherin (**Fig. 3e**; ref. (Borghi et al., 2012)). This showed a gradual increase in FRET ratios corresponding to a reduction in tension on E-cadherin as monolayer density increases above 3000 cells / mm^2^ and plateauing at a density of ~4000 cells / mm^2^ (**Figs. 3f, S2c**). Importantly, this density-dependent increase in FRET level was not observed in a force-insensitive construct that is not linked to the actin cytoskeleton as it lacks part of the E-cadherin cytosolic tail (E-cadherin TsMod ΔCyto) (**Figs. 3f, S2c**). Thus, our data show that prolongation of G2-phases at high monolayer density coincides with a decrease in tensile forces on E-cadherin adhesions between neighboring cells.

**Figure 3.**
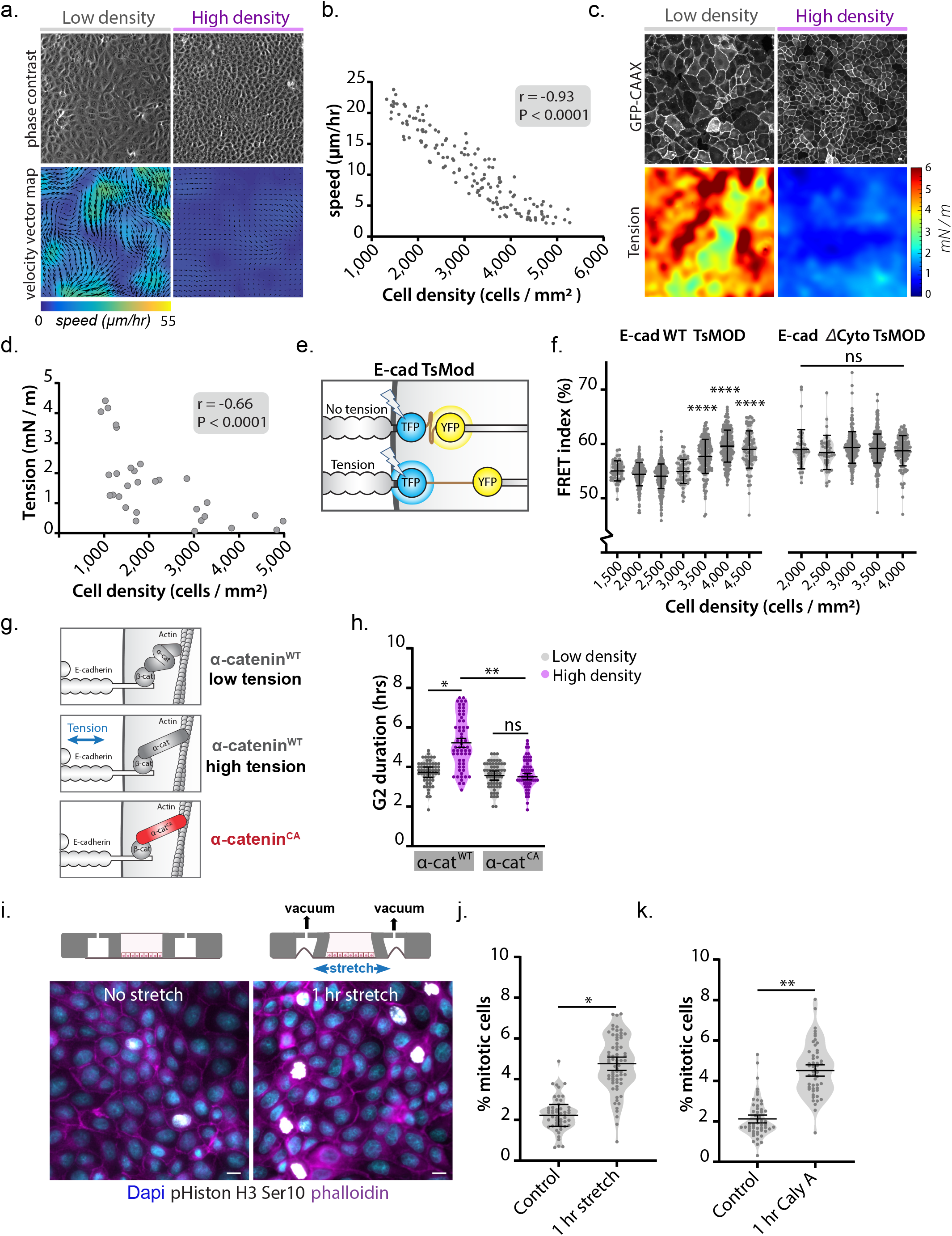
Density-dependent G2/M regulation requires mechanotransduction through E-cadherin adhesions. **(a)** Phase contrast images of MDCK monolayers grown at low and high density with corresponding vector magnitude heat maps of cell motility quantified by particle image velocimetry (PIV) analysis. The direction and length of the arrows indicate the direction and speed of motility. **(b)** Quantification of cell motility (μm / hr) at various MDCK monolayer densities. n = 150; data were pooled from 3 independent experiments. r = −0.93, P < 0.0001; Pearson correlation. **(c)** Representative examples of MDCK monolayer expressing GFP-CAAX at low and high density with maps of monolayer tension (tr(σ) · h [mN/m]) (corresponding traction force maps are shown in Fig. S2a). Scale bars represent 10 μm. **(d)** Quantification of average normal monolayer tension (tr(σ) · h [mN/m]) at various MDCK monolayer densities. n = 30; data were pooled from 7 independent experiments. r = −0.66, P < 0.0001; Pearson correlation (corresponding traction forces are shown in Fig. S2b). **(e)** Schematic representation of the FRET-based E-cadherin tension sensor (E-cadherin TsMod), in which a mTFP1/YFP FRET pair separated by a flexible linker is placed in the E-cadherin cytosolic tail. As tension on E-cadherin adhesions increases, mTFP1 and YFP are pulled apart resulting in a decrease in FRET signal (mTFP1/YFP ratio). E-cadherin ΔCyto lacks part of the cytosolic tail and its linkage to the actin cytoskeleton, and is therefore insensitive to tensile forces. **(f)** Graphs showing the FRET index (%) of individual cell-cell contacts of cells expressing E-cadherin TsMod (left) or the E-cadherin-ΔCyto TsMod negative control sensor (right). The indicated cell densities each represent a small range of densities ± 250 cells / mm^2^. Data were pooled from two independent experiments. Black bars represent the mean and SD. ****p < 0.0001; ns = not significant; Mann-Whitney. **(g)** Schematic representation of conformationally active α-catenin (α-cateninCA; M319G, R326E α-catenin). Tension on the E-cadherin complex results in unfolding of α-catenin, which is mimicked by expression of the α-cateninCA mutant that is constitutively in an open conformation irrespective of changes in intercellular tension. **(h)** Quantification of the duration of G2 length (hrs), based on mTurquoise-SLBP(18-126) expression, in cells expressing either wildtype (WT) α-catenin-mCherry or constitutively open (CA) α-catenin-mCherry at low (grey) and high (magenta) monolayer density. n = 60 cells per condition. Data were pooled from 3 independent experiments. Black bars represent the mean and SD of the individual experiments. *P = 0.0116; **P = 0.0079; ns = not significant; paired t-test. **(i)** Representative image of MDCK monolayers grown at high density, with or without application of 18% uniaxial stretch (1 hr), in which mitotic cells are visualized by immunostaining for phospho-Histon H3 Ser10 together with phalloidin and Dapi. Scale bars represent 10 μm. **(j)** Quantification of the percentage of mitotic cells in MDCK monolayers grown at high density with or without application of 18% uniaxial stretch for 1 hr. n > 50 monolayer regions per condition. Data were pooled from 3 independent experiments. Black bars represent the mean and SD of the individual experiments. *P = 0.0229; paired t-test. **(k)** Quantification of the percentage of mitotic cells in MDCK monolayers grown at high density with or without 1 hr treatment with the myosin phosphatase inhibitor Calyculin A (10 ng/ml). n > 50 monolayer regions per condition. Data were pooled from 3 independent experiments. Black bars represent the mean and SD of the individual experiments. **P = 0.0023; paired t-test.

To test if changes in tension on E-cadherin adhesions are responsible for the density-dependent regulation of G2 length, we aimed to manipulate the mechanism by which E-cadherin adhesions transduce forces. Tensile forces induce a conformational opening of the cadherin-complex component α-catenin, which ultimately results in the force-sensitive recruitment of additional proteins to the cadherin complex that can establish downstream signaling (Rauskolb et al., 2014; Yonemura et al., 2010). To render cadherin adhesions insensitive to fluctuations in intercellular forces, we expressed an α-catenin mutant that is constitutively in an open conformation, mimicking its tensed state irrespective of the level of intercellular forces (M319G, R326E α-catenin; α-catenin^CA^; **Figs. 3g, S2d**) (Maki et al., 2016; Matsuzawa et al., 2018). At low density, cells expressing GFP-tagged wildtype α-catenin (α-catenin^WT^) and α-catenin^CA^ showed similar G2 lengths (**Fig. 3h**). However, in contrast to the prolongation of G2 length at high density in α-catenin^WT^ cells, G2 length was not prolonged in α-cateninCA cells at high density (**Fig. 3h**). These data indicate that the increase in G2 duration at high density results from a decrease in mechanical tension that is sensed and transduced by E-cadherin-mediated cell-cell adhesions.

It has previously been established that levels of intercellular forces are elevated during epithelial expansion following removal of a physical constraint (Serra-Picamal et al., 2012). As we demonstrate that G2/M progression is regulated by intercellular forces transduced by E-cadherin adhesions, the rapid mitotic response during initiation of expansion of dense epithelia is thus presumably explained by force-dependent regulation of G2/M progression. To directly test whether an acute increase of intercellular forces can trigger G2 cells to enter mitosis, we artificially increased mechanical tension by application of uniaxial stretch to high density monolayers (**Fig. 3i**). We observed a rapid burst of mitotic events after 1 hour of mechanical stretch application (**Figs. 3i, 3j**), similar to the response during epithelial expansion following stencil release (**Figs. 1d, 1e**) and in line with recent findings in epithelial cultures and *Xenopus* embryos (Gudipaty et al., 2017; Nestor-Bergmann et al., 2019). Elevating forces in high density monolayers through myosin II-generated contractility by incubation with the myosin phosphatase inhibitor Calyculin A (Acharya et al., 2018) resulted in a comparable induction of mitotic events within hour (**Fig. 3k**). These data indicate that increasing mechanical tension relieves the G2 prolongation in dense epithelia and triggers rapid mitotic entry.

### An E-cadherin mechano-response controls levels of Wee1

To establish how E-cadherin-mediated transduction of intercellular forces regulates G2 length, we aimed to identify the molecular components downstream of E-cadherin that control the density-dependent G2/M transition. We analyzed potential changes in expression patterns of well-known regulators of G2/M progression between low and high density monolayers by Western Blot analysis, and observed in high density monolayers a strong upregulation of Wee1 (**Fig. 4a**); a central kinase in the regulation of G2/M transition following DNA damage (Hochegger et al., 2008; Lindqvist et al., 2009). Analyses of Wee1 by immunostaining validated the increase in Wee1 levels at high density compared to low density monolayers, with Wee1 accumulating both in the nucleus and cytosol (**Figs. 4b, 4c**). Importantly, expression of α-catenin^CA^ abrogated this Wee1 protein upregulation in dense monolayers, indicating that force transduction by the cadherin complex is essential for the density-dependent regulation of Wee1 (**Figs. 4d, S3a**).

**Figure 4.**
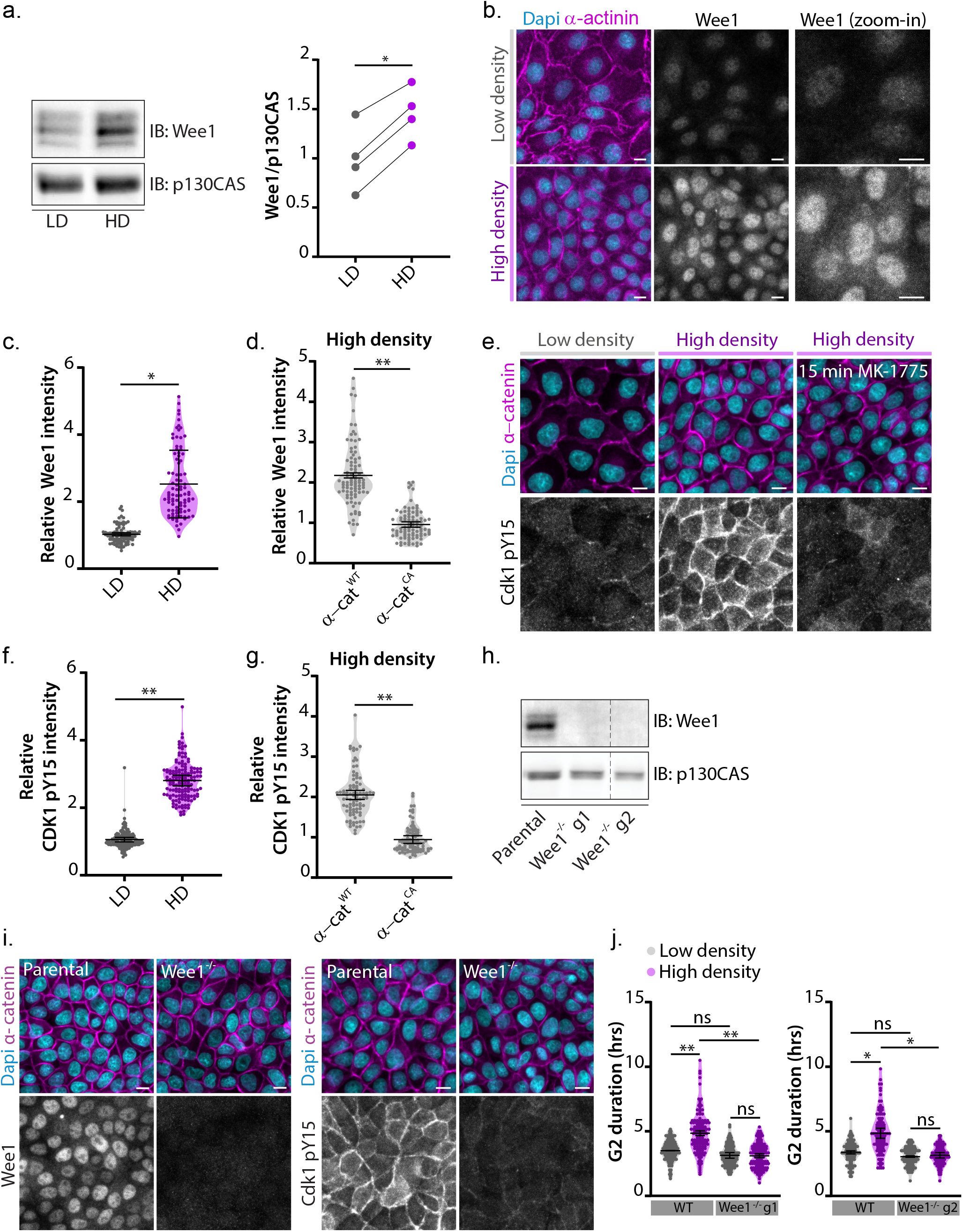
An E-cadherin mechanoresponse establishes density-dependent regulation of Wee1 levels. **(a)** Left: Western blot of lysates from MDCK cells grown at low (LD) and high (HD) monolayer density probed for Wee1 and p130CAS. Right: quantification of the ratio of Wee1 versus p130CAS of 4 independent experiments. *P = 0.0143; ratio paired t-test. **(b)** MDCK monolayers grown at low and high density and immunostained for Wee1 and α-actinin, together with Dapi. **(c)** Quantification of Wee1 immunofluorescence intensity per cell (normalized to the level of α-actinin) in MDCK cells grown at low and high monolayer density. n = 90 cells per condition. Data were pooled from 3 independent experiments. Black bars represent the mean and SD of the individual experiments. *P = 0.048; ratio paired t-test. **(d)** Quantification of Wee1 immunofluorescence intensity per cell (normalized to the level of β-catenin intensity) in MDCK cells expressing either wildtype (WT) α-catenin-mCherry or constitutively open (CA) α-catenin-mCherry. n = 90 cells per condition. Data were pooled from 3 independent experiments. Black bars represent the mean and SD of the individual experiments. **P = 0.0014; ratio paired t-test. **(e)** Immunostainings for Cdk1 pY15 and α-catenin, together with Dapi, in MDCK monolayers grown at low and high density, and at high density upon 15 min treatment with the Wee1 inhibitor MK-1775 (500 nM). **(f)** Quantification of Cdk1 pY15 immunofluorescence intensity per cell (normalized to the level of α-catenin intensity) in MDCK cells grown at low and high monolayer density. n = 150 cells per condition. Data were pooled from 3 independent experiments. Black bars represent the mean and SD of the individual experiments. **P = 0.0031; ratio paired t-test. **(g)** Quantification of Cdk1 pY15 immunofluorescence intensity per cell (normalized to the level of β-catenin intensity) in MDCK cells cultured at high density and expressing either wildtype (WT) α-catenin-mCherry or constitutively open (CA) α-catenin-mCherry. n = 90 cells per condition. Data were pooled from 3 independent experiments. Black bars represent the mean and SD of the individual experiments. **P = 0.0013; ratio paired t-test. **(h)** Western blot of lysates from parental and two Wee1^−/−^ MDCK clones (with independent guide sequences) probed for Wee1 and p130CAS. Dotted line indicates the region of the Western Blot that has been cropped out. **(i)** Immunostaining of parental and Wee1^−/−^ MDCK cells for Wee1 (left) and Cdk1 pY15 (right), together with α-catenin and Dapi. **(j)** Quantification of the duration of G2-phase, based on expression of mTurquoise-SLBP(18-126) in parental MDCK cells and the two independent Wee1^−/−^ clones grown at low and high monolayer density. n = 120 cells per condition. Data were pooled from 3 independent experiments. Black bars represent the mean and SD of the individual experiments. Left graph: **p = 0.0071 (low density WT vs. high density WT), **P = 0.0024 (high density WT vs. high density Wee1^−/−^ guide 1), right graph: *P = 0.0142 (low density WT vs. high density WT), *P = 0.0168 (high density WT vs. high density Wee1^−/−^ guide 2), ns = not significant; paired t-test. All scale bars represent 10 μm.

Wee1 regulates Cdk1 activity through an inhibitory phosphorylation of its Y15 residue (Parker et al., 1992). To test if this inactive state of Cdk1 is induced in high density monolayers, we immunostained epithelial monolayers with an antibody selectively recognizing pY15-phosphorylated Cdk1. This showed very low levels of Cdk1 pY15 at low monolayer density, which became strongly increased in dense monolayers (**Figs. 4e, 4f**). Because Cdk1 pY15 levels were elevated in all cells of the high density monolayer (**Figs. 4e, 4f**), this indicates that the negative regulation of Cdk1 occurs independently of the cell cycle phase in which cells reside. Treatment with the selective Wee1 inhibitor MK-1775 resulted in nearly complete disappearance of Cdk1 pY15 immunostaining, demonstrating that Y15 phosphorylation of Cdk1 in dense MDCK monolayers is predominantly mediated by Wee1 (**Fig. 4e**). Importantly, the increase in Cdk1 pY15 levels at high density was abolished by expression of α-catenin^CA^ (**Figs. 4g, S3b**). Altogether, our data indicate that Wee1 levels and concomitant inhibitory Cdk1 phosphorylation increase with monolayer density in a manner dependent on E-cadherin-mediated force transduction.

Of note, we observed that pY15-phosphorylated Cdk1 was not only present in the nucleus and cytosol, but also prominently resided at cell-cell contacts (**Fig. 4e**). To validate this junctional localization, we performed a transient CRISPR/CAS9-mediated knockout of Cdk1, which abolished the immunofluorescence signal of both total and Y15-phosphorylated Cdk1 in the nucleus and cytosol as well as at cell-cell junctions (**Figs. S3c – S3f**). Given that both the junctional and cytosolic pools of pY15-phosphorylated Cdk1 increase with cell density, potentially one or both of these pools may contribute to the density-dependent prolongation of G2-phase.

### Wee1 accumulation is required for G2 prolongation at high cell density

Having established the elevation of Wee1 levels and downstream Cdk1 Y15 phosphorylation in dense monolayers, we next investigated whether this is responsible for the G2 prolongation in these cells. To this end, we generated two Wee1 knockout (Wee1^−/−^) MDCK clones through CRISPR/Cas9-based gene editing using independent guide sequences (**Figs. 4h, 4i**), which also showed near complete loss of Cdk1 pY15 levels in dense monolayers (**Fig. 4i**). Of note, we did not observe any increase in DNA damage in Wee1^−/−^ cells under our culture conditions, as determined by yH2AX immunostaining (**Fig. S4a**). Next, we analyzed G2 duration in Wee1^−/−^ cells based on Clover-Geminin(1-110) and mTurquouise2-SLBP(18-126) expression. At low monolayer density, G2 length was comparable (~3 – 3.5 hours) between Wee1^−/−^ clones and parental cells (**Fig. 4j**), indicating that Wee1 is not essential per se for the regulation of G2/M progression in MDCK cells. However, the prolongation of G2-phase at high density was completely abolished in both Wee1^−/−^ clones (**Fig. 4j**). These data demonstrate that, besides its role in regulating G2/M progression upon DNA damage (Raleigh and O’Connell, 2000; Schmidt et al., 2017; Yarden et al., 2002), Wee1 plays a specific role in the control of the epithelial cell cycle by prolonging G2 length when monolayer density increases.

While expression of Wee1 is strongly increased in dense monolayers, Wee1 is not completely absent at low cell density (**Figs. 4a-4c**). As such, the complete loss of Wee1 in our knockout cells does not fully recapitulate the Wee1 status in low density monolayers. We also generated heterozygous Wee1 knockout MDCK cells (Wee1^−/+^) in which only one of the two Wee1 alleles is lost (**Fig. S4b**). At high monolayer density, the levels of Wee1 were substantially reduced in Wee1^−/+^ cells compared to wildtype cells, and remained more comparable to wildtype cells cultured at low density (**Figs. S4c, S4d**). This partial reduction of Wee1 was sufficient to abrogate the density-dependent G2 regulation, as both the elevated level of Cdk1 Y15 phosphorylation and the increase in G2 length at high monolayer density were lost in Wee^−/+^ cells (**Figs. S4e, S4f**). Altogether, these data demonstrate that prolongation of G2-phase at high monolayer density depends on the concurrent accumulation of Wee1.

### Tension-induced Wee1 degradation triggers mitotic entry

As we demonstrated that the rise of Wee1 levels is responsible for the prolongation of G2-phase at increasing cell density, we hypothesized that the induction of mitotic entry following elevation of intercellular forces requires downregulation of Wee1. To test this, we analyzed Wee1 levels in high density monolayers during epithelial expansion upon removal of the PDMS stencil (as in **Fig. 1c**). Within 1 hour following stencil removal, we observed a ~2-fold decrease in Wee1 levels (**Figs. 5a, 5b**). This rapid loss of Wee1 depends on its proteasomal degradation, as in the presence of the proteasome inhibitor MG132, Wee1 levels remained unaltered following stencil removal (**Figs. 5b, S5a**). Artificial increase in tension by application of mechanical stretch or Calyculin A treatment in high density monolayers resulted in a similar reduction of Wee1 levels (**Figs. 5c, S5b, S5c**), indicating that elevated mechanical tension triggers Wee1 degradation. In line with Wee1 being reduced, the levels of Cdk1 pY15 also declined within 1 hour after stencil removal (**Figs. 5d, S5d**) as well as upon application of mechanical stretch (**Figs. 5e, S5e**) or treatment with Calyculin A (**Fig. S5f**). Altogether, our data demonstrate that increasing mechanical tension triggers Wee1 degradation, thereby relieving the inhibitory state of Cdk1.

**Figure 5.**
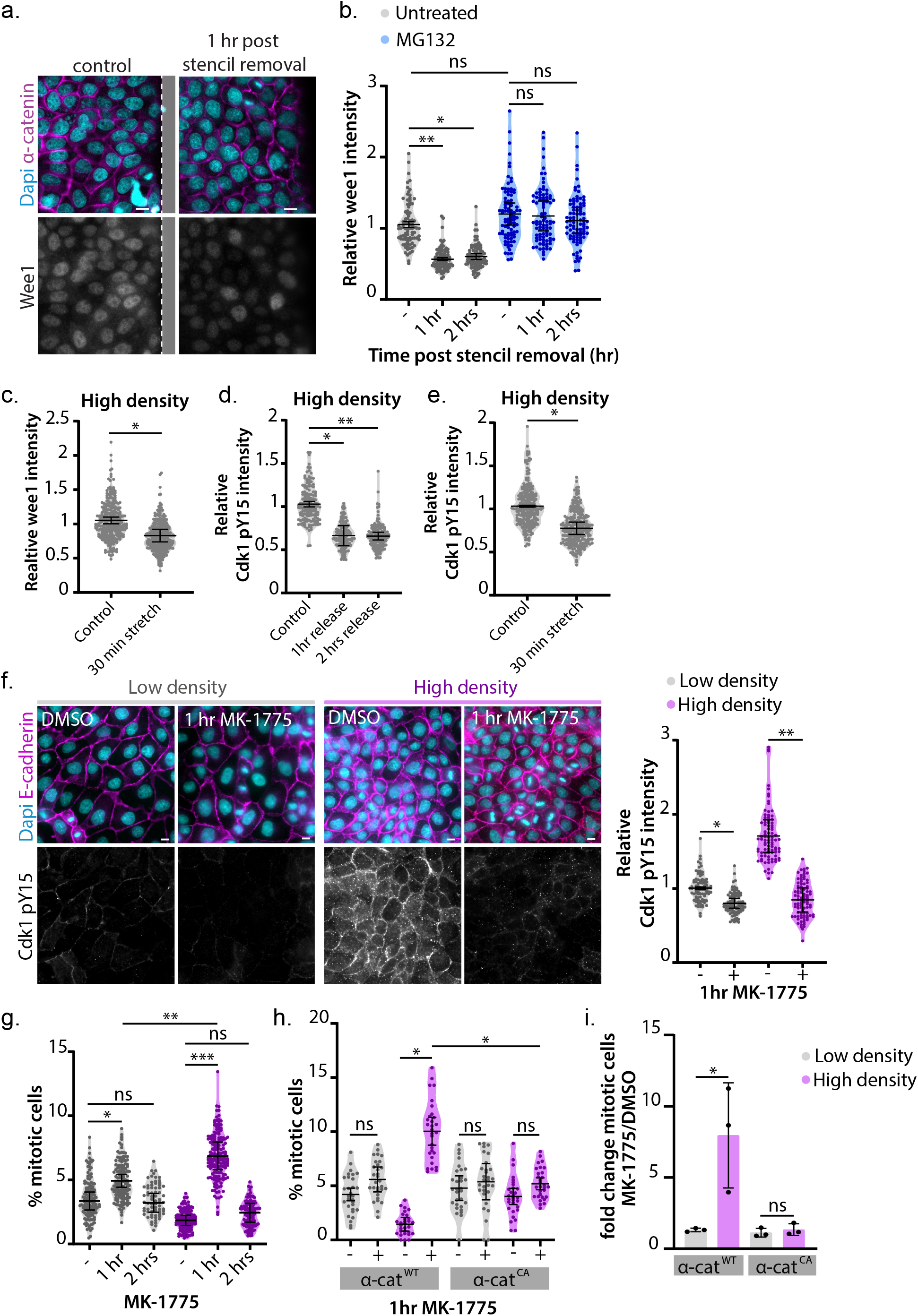
Tension-induced Wee1 degradation triggers mitotic entry. **(a)** Immunostainings of MDCK cells for Wee1, together with α-catenin and Dapi in an unperturbed high density monolayer and 1 hour after removal of a (polydimethylsiloxane) PDMS stencil (indicated in grey) to mimic epithelial expansion following wounding. **(b)** Quantification of Wee1 immunofluorescence intensity per cell (normalized to the level of α-catenin intensity) in unperturbed MDCK monolayers and 1hr or 2hr following removal of the PDMS stencil, either in absence (grey) or presence (blue) of the proteasome inhibitor MG132 (5 μm). MG132 was added 30 min before stencil release. n = 90 cells per condition. Data were pooled from 3 independent experiments. Black bars represent the mean and SD of the individual experiments. *P = 0.012; **P = 0.0013; ns = not significant; ratio paired t-test. **(c)** Quantification of Wee1 immunofluorescence intensity per cell (normalized to the level of α-catenin intensity) in MDCK monolayers with and without application of 18% uniaxial stretch (30 min). n = 290 cells per condition. Data were pooled from 4 independent experiments. Black bars represent the mean and SD of the individual experiments. *P < 0.027; ratio paired t-test. **(d)** Quantification of Cdk1 pY15 immunofluorescence intensity per cell (normalized to the level of α-catenin intensity in unperturbed MDCK monolayers and 1 hr or 2 hrs following removal of the PDMS stencil. n = 150 cells per condition. Data were pooled from 3 independent experiments. Black bars represent the mean and SD of the individual experiments.*P = 0.034; **P = 0.0049; ratio paired t-test. **(e)** Quantification of Cdk1 pY15 immunofluorescence intensity per cell (normalized to the level of α-catenin intensity) in MDCK monolayers with and without application of 18% uniaxial stretch (30 min). n = 240 cells per condition. Data were pooled from 3 independent experiments. Black bars represent the mean and SD of the individual experiments. *P = 0.0397; ratio paired t-test. **(f)** Left: Immunostainings for Cdk1 pY15, together with α-catenin and Dapi, of MDCK monolayers grown at low (left) and high (right) density, either in the absence or presence of the Wee1 inhibitor MK-1775 (1hr; 500 nM). Right: Quantification of Cdk1 pY15 immunofluorescence intensity per cell (normalized to the level of E-cadherin intensity) in MDCK monolayers grown at low (grey) or high (magenta) density, either in the absence or presence of the Wee1 inhibitor MK-1775 (1hr; 500 nM). n= 90 cells per condition. Data were pooled from 3 independent experiments. Black bars represent the mean and SD of the individual experiments.*P = 0.032; **P = 0.0093; ratio paired t-test. **(g)** Quantification of the percentage of mitotic cells in parental MDCK monolayers grown at low (grey) and high (magenta) density, treated either with DMSO or 1hr or 2 hrs with MK-1775 (500 nM). n > 69 fields of view per condition. Data were pooled from 4 independent experiments (3 independent experiments for the 2 hr timepoint). Black bars represent the mean and SD of the individual experiments. *P = 0.021; **P = 0.007; ***P = 0.0009, ns = not significant; paired t-test. **(h)** Quantification of the percentage of mitotic cells in MDCK cells expressing wildtype (WT) α-catenin-mCherry or constitutively open (CA) α-catenin-mCherry, at low (grey) and high (magenta) density, and with or without 1hr treatment with MK-1775 (500 nM). n= 30 monolayer regions per condition. Data were pooled from 3 independent experiments. Black bars represent the mean and SD of the individual experiments. *P = 0.0161 (high density WT untreated vs high density WT MK-1775 treated); *P = 0.0136 (high density WT MK-1775 treated vs. high density CA MK-1775 treated), ns = not significant; paired t-test. **(i)** Bar graph showing the fold-change in the percentage of mitotic cells following 1 hr treatment with MK-1175 (500 nM) in MDCK monolayers expressing either wildtype (WT) α-catenin-mCherry or constitutively open (CA) α-catenin-mCherry and grown at low (grey) and high (magenta) monolayer density (from data shown in Fig. 5h). The black dots represent the mean of each experiment and the black bars represent the SD. *P = 0.022; ns = not significant; ratio paired t-test. All scale bars represent 10 μm.

Finally, we directly tested if the downregulation of Wee1 and accompanying loss of Cdk1 pY15 are responsible for the increase in mitotic events downstream of increased mechanical tension in dense monolayers. To address this, we reduced Wee1 activity with its selective inhibitor MK-1775, leaving levels of mechanical tension unaltered. In dense monolayers, we observed a substantial reduction in Cdk1 pY15 levels and a 3.7-fold increase in the percentage of mitotic events upon 1 hour of Wee1 inhibition (**Figs. 5f, 5g**). This burst in mitosis was transient, as the level of mitotic events was almost completely restored to normal after hours (**Fig. 5g**). In contrast to dense monolayers, low density monolayers only showed a minor, albeit significant, increase in mitosis within an hour (1.47-fold increase). Moreover, the total percentage of mitotic cells at 1hr after Wee1 inhibition remained significantly lower in these low density monolayers compared to the MK-1775-treated dense monolayers (**Fig. 5g**). Similarly, in dense monolayers of cells expressing α-catenin^CA^, which lack a population of cells with G2 prolongation (**Fig. 3h**), inhibition of Wee1 was much less effective in increasing mitotic events than in dense α-catenin^WT^ monolayers (**Figs. 5h, 5i**). These data indicate that specific reduction of Wee1 activity, which occurs upon elevation of mechanical tension, is sufficient to relieve the density-dependent prolongation of G2-phase and triggers rapid mitotic entry. Taken together, our data show that Wee1 protein levels are mechanically controlled by intercellular forces transduced by E-cadherin adhesion to establish cell density-dependent regulation of G2/M progression.

## Discussion

Our work has uncovered that, in addition to conventional contact inhibition of proliferation that halts cells in G0/ G1, epithelial cells possess a mechanical checkpoint that serves to coordinate G2/M progression with local variations in cell density. Mechanistically, we show this involves mechanosensing by E-cadherin-based cell-cell adhesions and subsequent regulation of the Cdk1-inhibitory kinase Wee1 (**Fig. 6**). As local cell density increases, tension on E-cadherin adhesions is reduced, which results in the accumulation of Wee1 to maintain an inactive state of Cdk1. The density-dependent regulation of Wee1 and downstream Cdk1 occurs throughout the cell cycle, and once cells enter G2 is sufficient to temporarily halt cells in this phase of cell cycle. This establishes a pool of cells that can be readily triggered to enter mitosis once the inactive state of Cdk1 is relieved. Our data show that this release of Cdk1 inhibition is induced by the elevation of intercellular tension, for instance during epithelial expansion upon wounding, as this triggers rapid degradation of Wee1 and subsequent mitotic entry. Our data thus demonstrate that mammalian cells are responsive to mechanical cues from their environment during G2-phase, which may provide a mechanism to ensure epithelial integrity by rapid increase in cell number without a time-delay needed for progression through the entire cell cycle.

**Figure 6.**
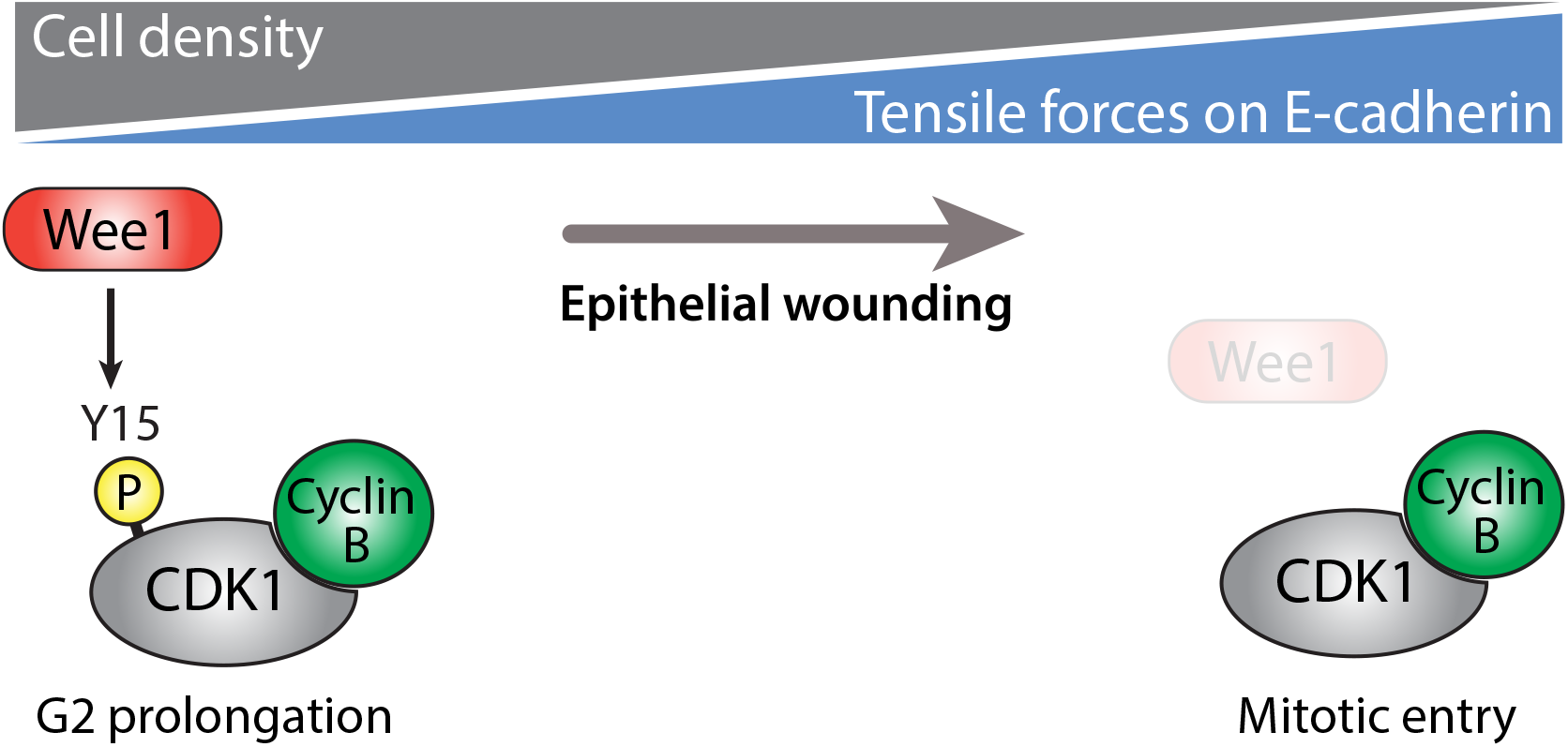
Model of the force-dependent regulation of Wee1-Cdk1 by E-cadherin. As epithelial monolayers increase in cell density, intercellular tensile forces are gradually reduced. This reduction in tension is sensed by E-cadherin-mediated cell-cell adhesions, which induce accumulation of the kinase Wee1 that phosphorylates the mitotic kinase Cdk1 on its Y15 residue, thereby inhibiting Cdk1 activity irrespective of the cell cycle phase in which cells reside. Once cells enter G2-phase, this inhibitory state of Cdk1 is sufficient to maintain cells in G2-phase for a prolonged period of time, ultimately resulting in an increased population of G2 cells at high monolayer density. These G2 cells can be rapidly triggered to divide upon an acute increase in tensile force, for instance during epithelial expansion following wounding, as this triggers proteasome-dependent degradation of Wee1 and subsequent loss of the inhibitory Y15 phosphorylation of Cdk1.

While it has long been thought that cells become irresponsive to extracellular cues following G1/S progression (Pardee, 1989), several studies in different model organisms support our findings indicating G2-phase as a key point of cell cycle control by extrinsic cues. As early as in the 1960’s and 1970’s, labeling studies performed in mouse and rat epithelia suggest the presence of a pool of cells that are blocked in G2-phase indefinitely, yet these cells maintain the capacity to divide in response to external stimuli, such as wounding or hormones (Gelfant, 1977; T. Pedersen, 1970). Recent studies in *Drosophila*, *Xenopus Laevis* and *C. elegans* tissues uncovered that populations of adult stem cells are halted in G2-phase and can be triggered to divide by growth factor and nutrient signaling (McKeown and Cline, 2019; Morris and Spradling, 2011; Otsuki and Brand, 2018; Seidel and Kimble, 2015; Zielke et al., 2014). Extrinsic control of G2/M progression may thus be a widely conserved element of the eukaryotic cell cycle. Our work, supported by recent findings of stretch-induced regulation of mitotic events (Gudipaty et al., 2017; Nestor-Bergmann et al., 2019), shows that this involves a key role for intercellular communication through mechanical forces. Our findings establish that extrinsic cues originating from cell density and concurrent levels in intercellular tension are major determinants in the control of G2 length and subsequent mitotic progression in mammalian epithelia. This is also in line with recent work demonstrating levels of intercellular tension as a predictor of cell cycle duration, including the combined length of S-G2-M phases (Uroz et al., 2018). The mechanical G2 checkpoint may serve to ensure precise coordination of cell division with variations in local cell density that occur, for instance, following wounding of the epithelium. Besides enabling an immediate increase in cell numbers following wounding, halting cells in G2 rather than cell cycle exit may have additional advantages as it allows for high fidelity homologous recombination-mediated repair in response to DNA damage, thereby preserving the integrity of the genome.

Besides implications for epithelial homeostasis, the mechanical G2 checkpoint might also be involved in the control of cell division in tissue development. During embryogenesis, cells are temporarily blocked in G2 in order to coordinate cell divisions with tissue growth, differentiation and morphogenetic movements (Bouldin and Kimelman, 2014; Grosshans and Wieschaus, 2000; Leise and Mueller, 2002; Mata et al., 2000; Murakami et al., 2004; Ogura et al., 2011; Seher and Leptin, 2000). As morphogenesis is inherently a mechanical process, variations in intercellular tension that occur during morphogenetic movements could potentially contribute to this cell cycle regulation.

Our data identify that Wee1 is not only a key component of DNA-damage induced cell cycle arrest, but also part of a mechanical G2 checkpoint that is regulated by E-cadherin mechanosensing and the force-induced conformational opening of α-catenin (**Fig. 3**) The regulation of the Hippo pathway by intercellular forces that underlies the mechanical regulation of G0/G1 contact inhibition is also established by force-induced opening of α-catenin and consequent binding to Ajuba/Zyxin family proteins, which then sequester the kinase LATS in an inactive state at AJs (Alégot et al., 2019; Dutta et al., 2018; Ibar et al., 2018; Rauskolb et al., 2014). Whether the conformational opening of α-catenin directly establishes Wee1 regulation, or this involves indirect signaling (e.g. through modulation of the actin cytoskeleton), remains an important question for future investigation. Our data indicate that intercellular forces regulate Wee1 at the level of its proteasomal degradation (**Fig. 5b**), which is known to be controlled by the ubiquitin ligases β-trcp and Trigger of mitotic entry-1 (Tome-1) (Smith et al., 2007). Tome-1 has previously been identified as a potential interactor of the E-cadherin complex (Guo et al., 2014), suggesting a putative link between E-cadherin adhesions and Wee1 degradation. Intriguingly, in budding yeast Wee1 regulation couples the timing of mitotic entry to cell growth, inducing division only once cells have achieved their appropriate size (Allard et al., 2017). This suggests an evolutionary conserved role for Wee1 in sensing physical cues and coordinating cell divisions with cell/tissue size.

Although our data clearly demonstrate a role for E-cadherin adhesions and Wee1 in mediating the density-dependent G2/M progression, importantly, this does not exclude the contribution of other mechanotransduction mechanisms. As such, it was recently shown that application of mechanical stretch triggers mitotic entry through activation of the calcium-channel Piezo1 (Gudipaty et al., 2017). This function of Piezo has been linked to regulation of cyclin B transcription (Gudipaty et al., 2017), and we find that direct activation of Piezo1 and downstream calcium influx does not affect Wee1 levels in MDCK cells (**Fig. S6**). This suggests E-cadherin and Piezo1 may act in parallel to control force-dependent G2/M progression. Future studies may shed light on the interplay between different mechanosensing complexes in the regulation of G2/M progression in epithelial cells.

We demonstrate that the prolongation of G2-phase at high density specifically depends on accumulation of Wee1 (**Figs. 4j, S4f**), and moreover, that inhibition of Wee1 is sufficient to drive these G2-halted cells into mitosis (**Figs. 5g-5i**). Nonetheless, as the inhibitory phosphorylation of Cdk1 by Wee1 is counteracted by Cdc25 phosphatases, the activation of Cdk1-Cyclin B ultimately depends on the balance between Wee1 and Cdc25 activity (Lindqvist et al., 2009). As such, it will be interesting to explore if epithelial wounding and fluctuations in cell density specifically control Wee1 levels, or also impact Cdc25.

We observed that in epithelial cells Cdk1 and its pY15-phosphorylated state are not only present in the nucleus and cytosol, but also reside at cell-cell contacts (**Figs. 4e, S3c**). This is in line with recent findings in *Xenopus* embryos, where a large fraction of Cdk1 and the pY15-phosphorylated pool was shown to localize to cell-cell junctions (Sandquist et al., 2018). The density-dependent accumulation of Cdk1 pY15 at cell-cell junctions was abolished following Wee1 inhibition (**Fig. 4e**), implying that the cytosolic Wee1 pool is able to phosphorylate Cdk1 at junctions. Our data indicate that both the junctional and cytosolic Cdk1 pools undergo density-dependent regulation of the inhibitory phosphorylation by Wee1 (**Fig. 4e**), yet it remains to be determined which fraction of Cdk1 is essential for the density-dependent regulation of G2/M progression. It has been proposed that Cdk1-Cyclin B is first activated in the cytoplasm prior to mitotic entry and then stimulates its own transport to the nucleus by regulating the nuclear transport machinery (Gavet and Pines, 2010). Whether junctional Cdk1 can translocate to the nucleus, or whether it influences cytosolic Cdk1 through one of the described positive feedback mechanisms by which Cdk1 promotes its own activity (Lindqvist et al., 2009), remains to be determined. Cdk1 may also perform specific functions at cell-cell junctions. For instance, in *Xenopus* embryos activation of the junctional Cdk1 pool during metaphase is implicated in coupling mitotic spindle dynamics to anaphase onset (Sandquist et al., 2018). One alluring hypothesis is that regulation of the junctional Cdk1 pool may modulate cell-cell adhesion complexes throughout the cell cycle, for instance to allow their remodeling before mitotic entry to enable mitotic rounding, analogous to the regulation of focal adhesion dynamics by Cdk1 (Jones et al., 2018). Various junctional- and actin cytoskeleton components were identified as potential Cdk1 substrates in an unbiased quantitative phosphoproteomic analysis in mitotic HeLa cells (Petrone et al., 2016), and it will be interesting to test their potential contribution to cell cycle-dependent junctional remodeling.

## Materials and methods

### Antibodies and reagents

The following commercial antibodies were used at the indicated concentrations for Western blot (WB) and immunofluorescence (IF): α-tubulin (DM1A, Sigma; T6199, 1:1000 IF), Ki67 Abcam; cat# ab15580; 1:1000 IF), E-cadherin (DECMA-1, GeneTex; GTX11512, 1:1000 IF), αE-catenin (Enzo; ALX-804-101-C100, 1:500 IF), Phospho-Histone H3 Ser10 (3H10, Millipore; 05-806, 1:500 IF), Wee1 (D10D2, Cell signaling; 13084, 1:500 IF, 1:1000 WB), α-actinin (BM-75.2, Sigma; A5044, 1:500 IF), phospho-Cdk1 Tyr15 (Cell signaling; 9111, 1:125 IF), vinculin (hVIN-1, Sigma; V9131, 1:250 IF), Cdk1 (PSTAIR; Millipore; 06-923; 1:500), Cdk1 (POH1, Cell signaling; 9116, 1:250 IF), p130CAS (BD Biosciences; 610271, 1:3000 WB), phospho-Histone H2A.X Ser139 (JBW301; Millipore; 05-636, 1:1000 IF).

The following reagents were used at the indicated concentrations: MK 1775 (Axon Medchem; 1494, 500 nM), Calyculin A (Enzo; BML-EI192-0100, 10 ng/ml), Thapsigargin (Sigma; T9033, 1 μM), Yoda (Tocris; 5586, 10 μM).

### Plasmids

The following plasmids are described elsewhere: pLL3. 7m-Clover-Geminin(1-110)-IRES-mKO 2-Cdt(30-120) and pLL3.7m-mTurquoise2-SLBP(18-126)-IRES-H1-mMaroon1 (FUCCI4; (Bajar et al., 2016), Tol2-hEF1α-H2B TagRFP-2A-GCaMP3 (Tian et al., 2009) (kind gift from Bas Ponsioen, University Medical Center Utrecht, the Netherlands). The pInducer20 mCherry-αE-catenin plasmid was generated using In-Fusion cloning (inserting mCherry-αE-catenin [GI: 49935;(Oldenburg et al., 2015)] in a pInducer20 vector [Addgene; #44012]). Finally, the M319G, R326E αE-catenin (α-cateninCA; ref. (Matsuzawa et al., 2018)) mutant was created using PCR mutagenesis.

### Cell lines and culture

MDCK GII cells were cultured at 37 °C and 5% CO_2_ in low glucose DMEM containing 10% FBS, 1 g/L sodium bicarbonate, and penicillin/streptomycin. Live-cell imaging was performed with the same media formulations. MDCK E-cadherin TSMod and TSModΔCyto cell lines are previously described (Borghi et al., 2012), all other cell lines were generated using transfection reagent Lipofectamine 3000 (Thermo Fisher Scientific) or by lentiviral transductions and sorted by FACS to obtain monoclonal lines. Wee1 knockout lines were generated by expression of a SpCas9-GFP plasmid containing a Wee1-specific gRNA (guide 1: 5’-CACCGCGCGATGAGCTTCCTGAGC-3’, guide 2: 5’-CACCGAAGAGCCGCAGCTTGCGGA-3’). Knockout lines were verified by immunofluorescence and western blot analysis for endogenous Wee1. Cdk1 knockout cells were generated by transient expression of a SpCas9-GFP plasmid containing a Cdk1-specific gRNA (guide 1: 5’-AAACATGGAGTTGTGTATAAGGGTC-3’, guide 2: 5’-CACCGTTGTCTATTTCAGGTACCTA3’), and verified by Cdk1 immunostainings. For cells with doxycycline-induced α-catenin expression, cells were cultured in medium containing 2 μg/ml doxycycline for at least 24 hrs. All cell lines were regularly tested for the absence of mycoplasma.

### Live cell microscopy and analyses

For live cell imaging, cells were seeded on glass-bottom dishes (WillCo-dish^®^) pre-coated with Rat tail Collagen I (Corning), and imaged on a Zeiss Cell Observer equipped with Orca Flash 4.0 camera (Hamamatsu) using a 40× objective (NA = 1.1) (Fig. 2) or a 10x objective (NA = 0.3) (Fig. 3b), or on a Nikon Spinning Disc confocal microscope using a 20× objective (NA = 0.75) (Figs. 3h, 4j, S4f, S6b – d). Imaging was performed in temperature- and CO_2_-controlled incubators, using Zen image acquisition software and NIS-Elements software, respectively. For cell cycle analyses, MDCK cells expressing FUCCI4, or mTurquoise-SLBP(18-126) and Histon1-mMaroon to monitor only G2 length, were imaged with a time interval of 10 min. The length of each cell cycle phase was determined per cell based on changes in expression levels of the different cell cycle markers using ImageJ software (National Institutes of Health. The length of G0/G1 is defined as the time from completion of cell division (and concomitant start of Cdt1 expression) to the start of Cdt1 degradation; the length of S-phase is defined as the time from the start of Cdt1 degradation (and simultaneous increase in geminin expression) to the start of SLBP degradation; the length of G2-phase is defined as the time from the start of SLBP degradation to nuclear envelope breakdown. To monitor intracellular calcium levels, MDCK cells expressing the GCaMP3 calcium sensor were imaged with a time interval of 30 sec, before and after incubation with the indicated compounds. The GCaMP3 fluorescence signal (mean gray value) was measured over time in the cytosol of single cells using ImageJ software (National Institutes of Health).

### Immunofluorescence stainings and analyses

For immunofluorescence stainings, cells were fixed with 4% formaldehyde (SigmaAldrich), permeabilized with 0.2% Triton X-100 (SigmaAldrich), blocked in buffer containing 1% BSA (Sigma), 1% goat serum (Life Technologies), and 1% donkey serum (Jackson Immunoresearch), and incubated with the indicated primary and Alexa-conjugated secondary antibodies (Life technologies), together with Dapi (SigmaAldrich) where indicated. Cells were imaged on a Zeiss LSM880 scanning confocal microscope using a 40× objective (NA = 1.1), with the exception of Figs. 1a, 1d, 3i, 5f, S1a, S1f, S2d, S3e, S5d – f), which were imaged on a Zeiss Cell Observer equipped with Orca Flash 4.0 camera (Hamamatsu) camera using a 40× objective (NA = 1.1), using Zen image acquisition software.

The fluorescence intensities (mean gray value) of Wee1 and Cdk1 pY15 were measured in individual cells within the monolayer (with at least 10 cells analyzed per field of view) using ImageJ software (National Institutes of Health) and normalized to the intensity of co-stained control proteins as indicated. To compare results between different experiments, all values within one experiment were scaled to the median of the control condition of that experiment. Cells were imaged and included for analyses in an un-biased manner. Statistical analyses were performed on the means of independent experiments of the original (not re-scaled) data.

### Epithelial expansion upon stencil removal and stretch assays

To mimic epithelial wounding and induce epithelial expansion, cells were seeded at increasing densities within rectangular microwells that were cut using a computer-controlled razor writer (Cameo, Silhouette) in 250 μm thick silicone PDMS sheeting (Bisco HT-6240, Stockwell Elastomers). These silicone stencils were placed on top of a glass-bottom dish (WillCo-dish^®^) pre-coated with Rat tail Collagen I (Corning). Approximately 24 hrs after seeding, the PDMS membrane was carefully removed and cells were fixed at the indicated timepoints after removal of the PDMS membrane. Mitotic cells were visualized by immunostaining for phospho-Histon H3 Ser10.

For stretch assays, MDCK cells were plated on a PDMS silicone-based cell stretching device (previously described by (Hart et al., 2017). For this, PDMS elastomer and curing agent (Sylgard 182, Dow Corning) were mixed at 10:1 wt/wt ratio and poured over a 3D-printed mold and baked at 65 °C overnight. To smoothen the surface the device was stamped on a thin layer of uncured PDMS, followed by baking at 65 °C for 3 hrs. Subsequently, the device was bonded with the silicone membrane using plasma activation of the surface (PDC-32G, Harrick Plasma). Following coating with rat tail collagen type I (Corning), MDCK cells were seeded in the central channel. One day following plating, cells were stretched using a pneumatic valve to control vacuum pressure in the two neighboring side channels.

### PIV analysis

Cell motility within monolayers was assessed by Particle Image Velocimetry using the MATLAB add-on PIVLab (pivlab.blogspot.com) (Thielicke and Stamhuis, 2014). First, 16-bit phase contrast movies (time resolution 10 mins/frame, spatial resolution 59*106 pixels/ cm^2^) of MDCK cells at indicated monolayer densities were converted to 8-bit in ImageJ (v. 1.51). Images were loaded into PIVlab in the time-resolved image sequencing style and motility vectors were calculated using single pass direct cross correlation with an interrogation window of 24×24 pixels and a 50% overlap. Abnormally large vectors (vector magnitude > 5 pixels/ frame) were excluded from analysis. Vector magnitude heat maps were created using a parula color map (range 0-7 pixels/frame). Vector magnitude plots were created with GraphPad Prism (v9.1.0).

### Monolayer stress microscopy

Soft elastomeric silicone gels of 12.6 kPa stiffness were prepared adapting a previously published protocol (Bergert et al., 2016; Latorre et al., 2018). The two polydimethylsiloxane (PDMS) components CY52-276A and CY52-276B (Dow Corning Toray) were mixed at a 9:10 ratio, spin-coated on glass-bottom dishes (35-mm, no. 0 coverslip thickness, Mattek) and cured overnight. Soft PDMS gels were subsequently treated with (3-aminopropyl)triethoxysilane (APTES, Sigma-Aldrich, cat. no. A3648) diluted at 5% in absolute ethanol, rinsed 3 times with ethanol 96% and once with milli-Q water. Samples were then incubated for 5 min with a filtered (220 nm, Millex^®^ - GP) and sonicated solution of 200-nm-diameter red fluorescent carboxylate-modified beads (FluoSpheres, Invitrogen) in sodium tetraborate (3.8 mg/ml, Sigma-Aldrich), boric acid (5 mg/ml, Sigma-Aldrich) and 1-ethyl-3-(3-dimethylaminopropyl) carbodiimide (EDC, 0.1 mg/ml, Sigma-Aldrich). Gels were rinsed 3 times with milli-Q water and dried in the oven for 15 min at 60 °C. To prepare the PDMS gels for patterning after being coated with fluorescent beads, the surface of the gels was treated with a 1 mg/mL poly-lysine (PLL, Sigma) solution for 1h, then washed with HEPES buffer (pH=8.4). A solution of 50 mg/mL mPEG (MW 5,000) - succinimidyl valerate (SVA) (Laysan Bio) dissolved in the same HEPES buffer was applied to passivate the surface for 1h, and after washing with milli-Q water, micropatterns were generated by using a UV-activated mPEG-scission reaction spatially controlled by the system PRIMO (Alvéole) mounted on an inverted Nikon Eclipse Ti microscope. In the presence of a photo-initiator compound (PLPP, Alvéole), the antifouling properties of the PEGylated substrate are tuned by exposure to near-UV light (375 nm) and after illumination (800 mJ/mm^2^) through a 20x objective PLL is exposed. After rinsing with PBS, a mixture of rat tail type I collagen (8.33%, First Link (UK) Ltd.) and fluorescent Fibrinogen (647 nm, 1.67 %, Invitrogen) dilluted in PBS was incubated at room temperature for 5 min in order to coat the PEG-free PLL with the protein solution, and excess protein was washed out. Next, cells were plated (400 - 1200 cells / mm^2^) to obtain the desired densities after 24h, and the fluorescence signals of the beads and cells were acquired with an automatic inverted microscope (Nikon Eclipse Ti), with a spinning disk confocal unit (CSU-W1, Yokogawa), Zyla sCMOS camera (Andor, image size 2,048 × 2,048 pixels) controlled with Micro-Manager (Stuurman et al., 2007, 2010), using a 40x objective (NA 0.75, air) and thermal C0_2_ and humidity control. A tile of each island with 20% overlap was imaged using a motorized stage. The 2D displacement field of the top layer of the gel was measured using a custom-made PIV software in Matlab (MathWorks), by comparing the beads image of the deformed gel with a reference image taken after trypsinization. Traction forces were calculated using fineite-thickness Fourier-transform traction force microscopy (Trepat et al., 2009). Flat-field correction of the fluorescence signal of the individual tiles of the beads field was performed (Model, 2006; Model and Burkhardt, 2001). The corrected tiles were then stitched by using ImageJ’s Grid/Collection stitching plugin. Monolayer tension was calculated from the traction fields using Monolayer Stress Microscopy as described in (Serrano et al., 2019; Tambe et al., 2011, 2013), which was implemented as a custom-made software in Python 3 using NumPy, SciPy, Matplotlib, scikit-image, pandas, pyFFTW, opencv and cython.

### E-cadherin TsMod FRET imaging and analysis

E-cadherin TsMOD FRET experiments were performed as described in (Canever et al., 2020). MDCK cells expressing either E-cadherin TsMod or the force-insensitive control sensor E-cadherin TsModΔCyto were seeded at increasing densities on glass-bottom wells coated with Collagen IV. Spectral images were acquired on an LSM 780 confocal microscope (Zeiss) with a 63×/1.4NA oil-immersion objective. mTFP1 was excited by the 458-nm line of a 30-mW argon laser. Emission was sampled at a spectral resolution of 8.7 nm within the 470- to 600-nm range. Fluorescent images were analyzed in ImageJ using the Fiji distribution (http://fiji.sc/wiki/index.php/Fiji) and the publicly available PixFRET plugin [http://www.unil.ch/cig/page16989.html]. All channels were background-subtracted, Gaussian smoothed (radius = 1 pixel), and thresholded (above the first ~3–5% of the 12-bit range). The FRET index was computed as IEYFP/ (ImTFP + IEYFP) with I the intensity in the channel of maximum emission of the corresponding fluorophore. For quantification, the acceptor channel was used to segment the cell–cell contact regions, and index was averaged over the segmented cell–cell contacts.

## Acknowledgements

The laboratory of MG is supported by the Netherlands Organisation for Scientific Research (NWO; 016.Vidi.189.166, NWO gravitational program CancerGenomiCs.nl 024.001.028). The laboratory of NB is supported by the Centre national de la recherche scientifique (CNRS), the French National Research Agency (ANR) grants (ANR-17-CE13-0013, ANR-17-CE09-0019, ANR-18-CE13-0008), the Association pour la Recherche contre le Cancer (ARC-PJA22020060002255) and acknowledges the ImagoSeine facility, member of the France BioImaging infrastructure (ANR-10-INSB-04). HC is supported by La Ligue contre le Cancer (TAYS18872) and Fondation Recherche Médicale (FDT202001010843). We thank W. James Nelson (Stanford University), Blair Benham-Pyle (Stowers Institute) and members of our laboratories for helpful discussions, and Vinay Swaminathan (Lund University), Johan de Rooij (UMC Utrecht), Susanne Lens (UMC Utrecht), Geert Kops (Hubrecht Institute) and Ronja Houtekamer (UMC Utrecht) for critical reading of the manuscript.

## Supplemental figures

**Supplementary Figure 1.**
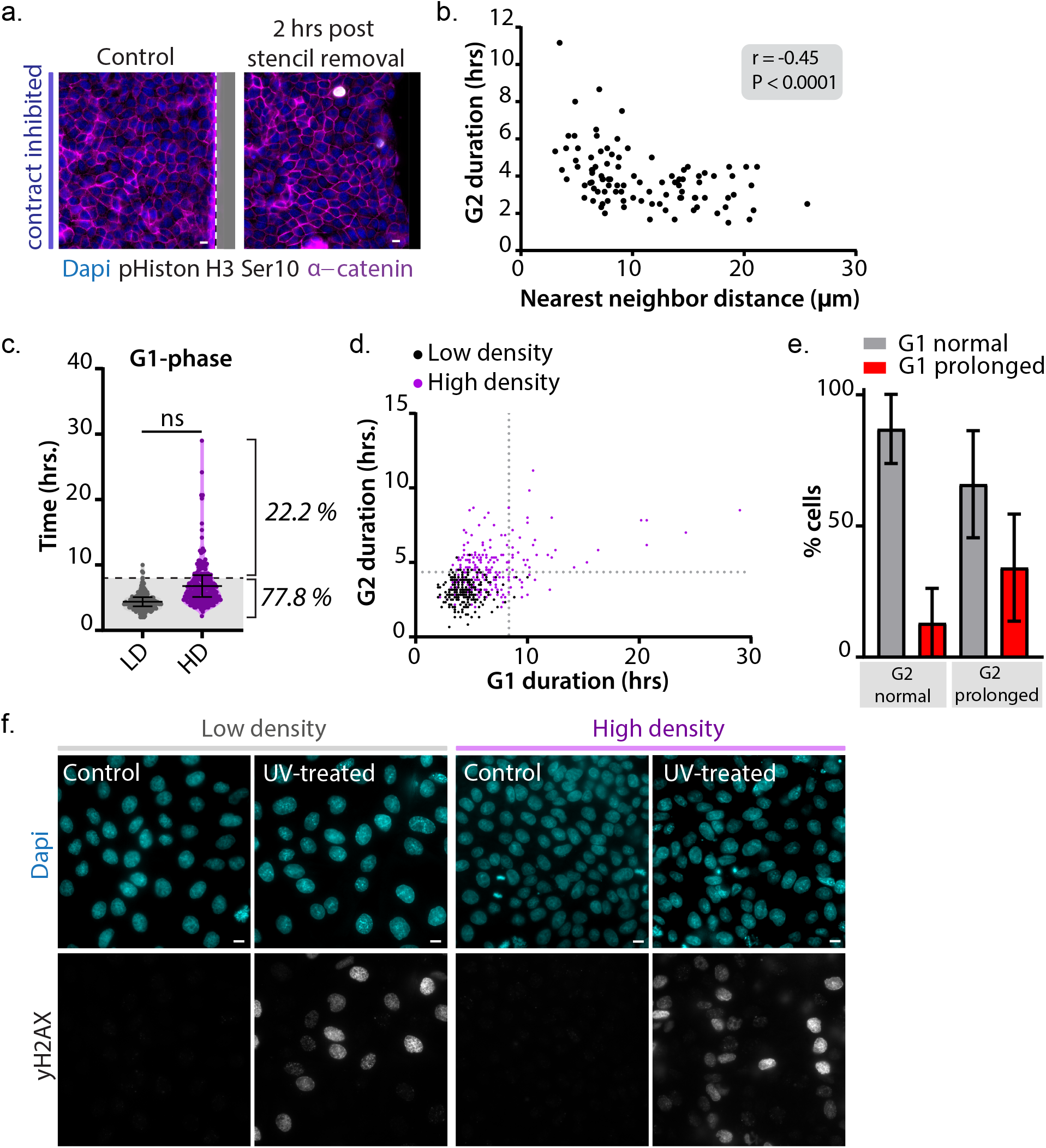
**(a)** Immunostaining of MDCK monolayers grown at contact-inhibited density (CIP; see Fig. 1b) without and 2 hours after induction of epithelial expansion by removal of the PDMS stencil, for phospho-Histon H3 Ser10 together with α-catenin and Dapi. The PDMS stencil is indicated in grey, and the dotted line indicates the border between the stencil and the monolayers of cells. **(b)** Quantification of the duration of G2-phase (hrs) of cells grown at various densities, correlated to the local nearest neighbor distance (NND; μm). Data shown is from one representative experiment. r = −0.45, P < 0.0001; Pearson correlation. **(c)** Quantification of the duration of G1-phase (hrs) in monolayers grown at low (grey) and high (magenta) density (same data as in fig. 2c), in which the 99% percentile of the duration of G1 in low density monolayers is indicated (8 hrs, dashed line), with 22.2 ± 16.07% of cells in high density monolayers showing a prolonged G1 duration longer than 8 hrs. Data were pooled from 3 independent experiments. Black bars represent the mean and SD of the individual experiments. ns = not significant; paired t-test. **(d)** Correlation of G1- and G2 length within individual cells at low (black) and high (magenta) monolayer density. The dotted lines indicate the 99 percentiles of G1-(x-axis; 8 hrs) and G2-(y-axis; 4.8 hrs) duration of cells at low density. n = 225 cells per condition. Data were pooled from 3 independent experiments. **(e)** Bar graph showing the correlation between G1 and G2 length in cells at high monolayer density. The majority of cells with a prolonged G2-phase (65.9 ± 20.4 %) did not show a similar prolongation of G0/G1 phase earlier in their cell cycle. Data were pooled from 3 independent experiments. Black bars represent the mean and SD of the individ ual experiments. **(f)** Immunostaining of MDCK monolayers grown at low (left) and high (right) density, with and without UV irradiation (20 J/m2) for the DNA damage marker yH2AX (visualizing DNA double strand breaks), together with Dapi. All scale bars represent 10 μm.

**Supplementary Figure 2.**
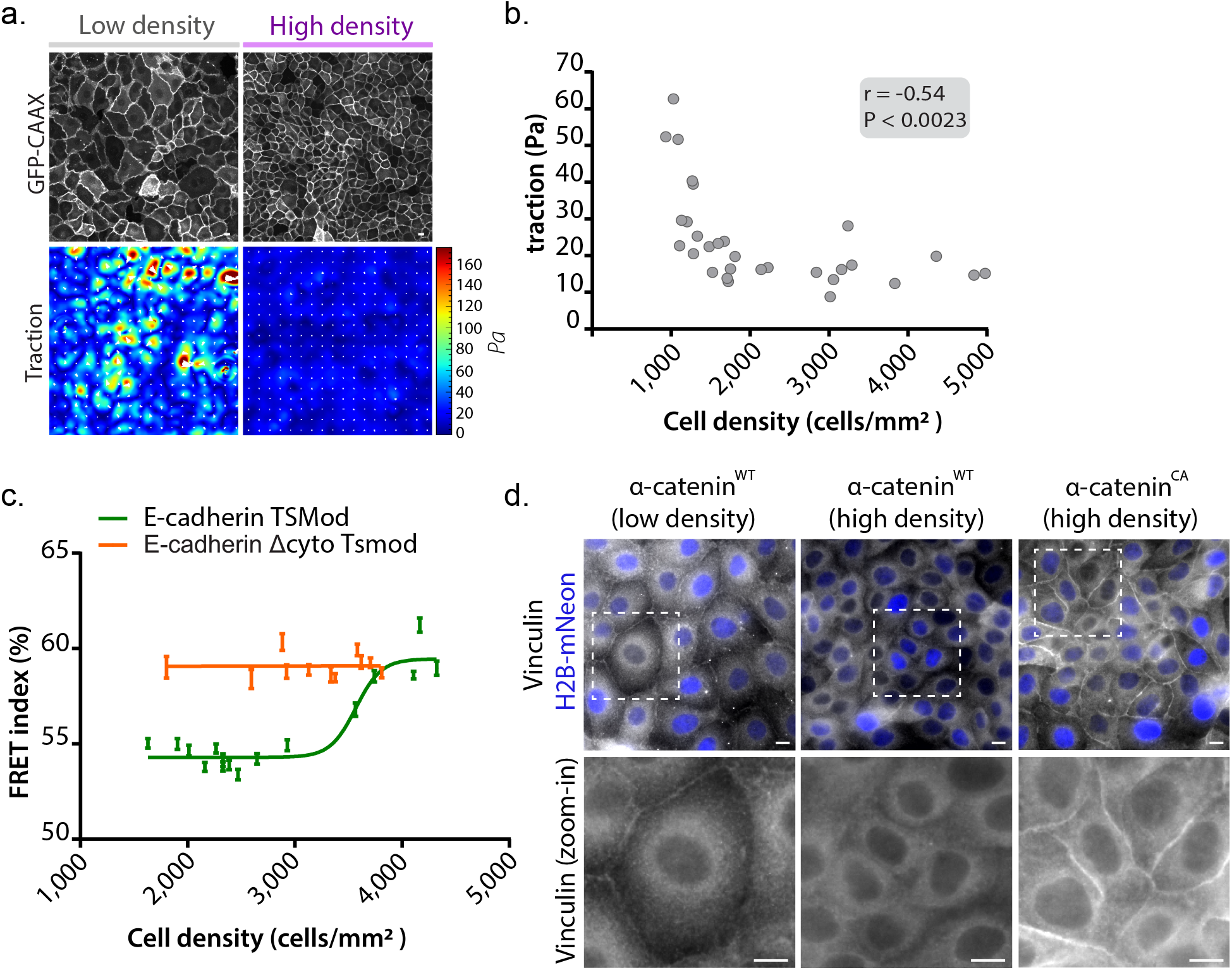
**(a)** Representative examples of MDCK monolayer expressing GFP-CAAX at low and high density with maps of traction forces (Pa). Scale bars represent 10 μm. **(b)** Quantification of average traction forces at various MDCK monolayer densities. n = 30; data were pooled from 7 independent experiments. r = −0.54, P < 0.0023; Pearson correlation. **(c)** Graph showing the correlation between the FRET index (%) and cell density (cells/mm^2^) of individual cell-cell contacts of cells expressing E-cadherin TsMod (green) or the E-cadherin E-ΔCyto TsMod negative control sensor (orange). Data were pooled from two independent experiments (same data as in fig. 3f). **(d)** Immunostainings of MDCK cells expressing either wildtype (WT) α-catenin-mCherry or constitutively open (CA) α-catenin-mCherry for endogenous vinculin. At low monolayer density tension on E-cadherin adhesions is high (Fig. 3f), resulting in α-catenin to unfold and recruitment of vinculin to adherens junctions. At high monolayer density, tension is low (Fig. 3f) and α-catenin adopts a closed conformation and consequently, vinculin is released from cell-cell contacts. In cells expressing the α-catenin^CA^ mutant, which is constitutively in the open conformation irrespective of changes in tension, vinculin is continuously bound to α-catenin and thus enriched at cell-cell contacts, even at high density. Scale bars represent 10 μm.

**Supplementary Figure 3.**
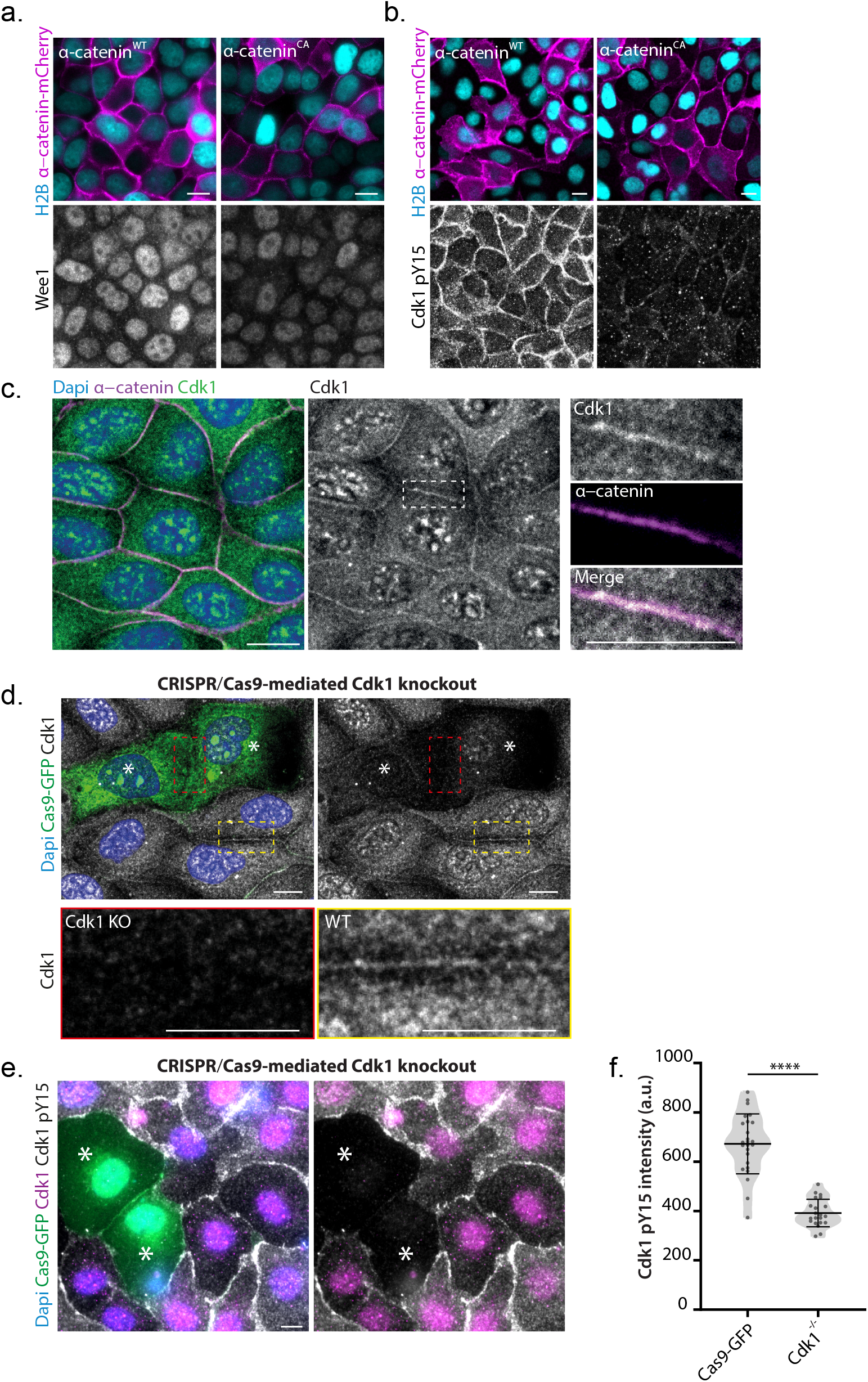
**(a)** Immunostainings of MDCK cells expressing either wildtype (WT) α-catenin-mCherry or constitutively open (CA) α-catenin-mCherry, together with H2B-mNeon, for Wee1. **(b)** Immunostainings of MDCK cells expressing either wildtype (WT) α-catenin-mCherry or constitutively open (CA) α-catenin-mCherry, together with H2B-mNeon, for Cdk1 pY15. **(c)** Immunostaining of MDCK cells for Cdk1 and α-catenin, together with Dapi. Inset shows an individual cell-cell contact. Note that cells were fixed with methanol in order to visualize the junctional pool of Cdk1. **(d)** Immunostainings of MDCK cells transiently expressing CRISPR/Cas9 with a Cdk1 targeting sequence, resulting in Cdk1 knockout, for Cdk1 together with Dapi. Note that cells were fixed with methanol in order to visualize the junctional pool of Cdk1. Cas9-GFP expressing cells were visualized by Cas9 immunostaining, Cas9-positive cells with Cdk1 knockout are indicated by white asterisks. Inset shows a cell-cell contact between two knockout cells (red), and between two wildtype cells (yellow). **(e)** Immunostainings of MDCK cells transiently expressing CRISPR/Cas9 with a Cdk1 targeting sequence, resulting in Cdk1 knockout, for Cdk1 and Cdk1 pY15. Cells expressing Cas9-GFP and showing Cdk1 (pY15) depletion are indicated by white asterisks. **(f)** Quantification of Cdk1 pY15 intensity in individual Cdk1^−/−^ cells (expressing Cas9-GFP with Cdk1 targeting sequence, and in which total Cdk1 levels were decreased) and in cells expressing a Cas9-GFP control vector. n = 25 cells (Cas9-GFP), n = 21 cells (Cdk1^−/−^). Data were derived from one representative experiment. Black bars represent the mean and SD. ****P < 0.0001; Mann-Whitney. All scale bars represent 10 μm.

**Supplementary Figure 4.**
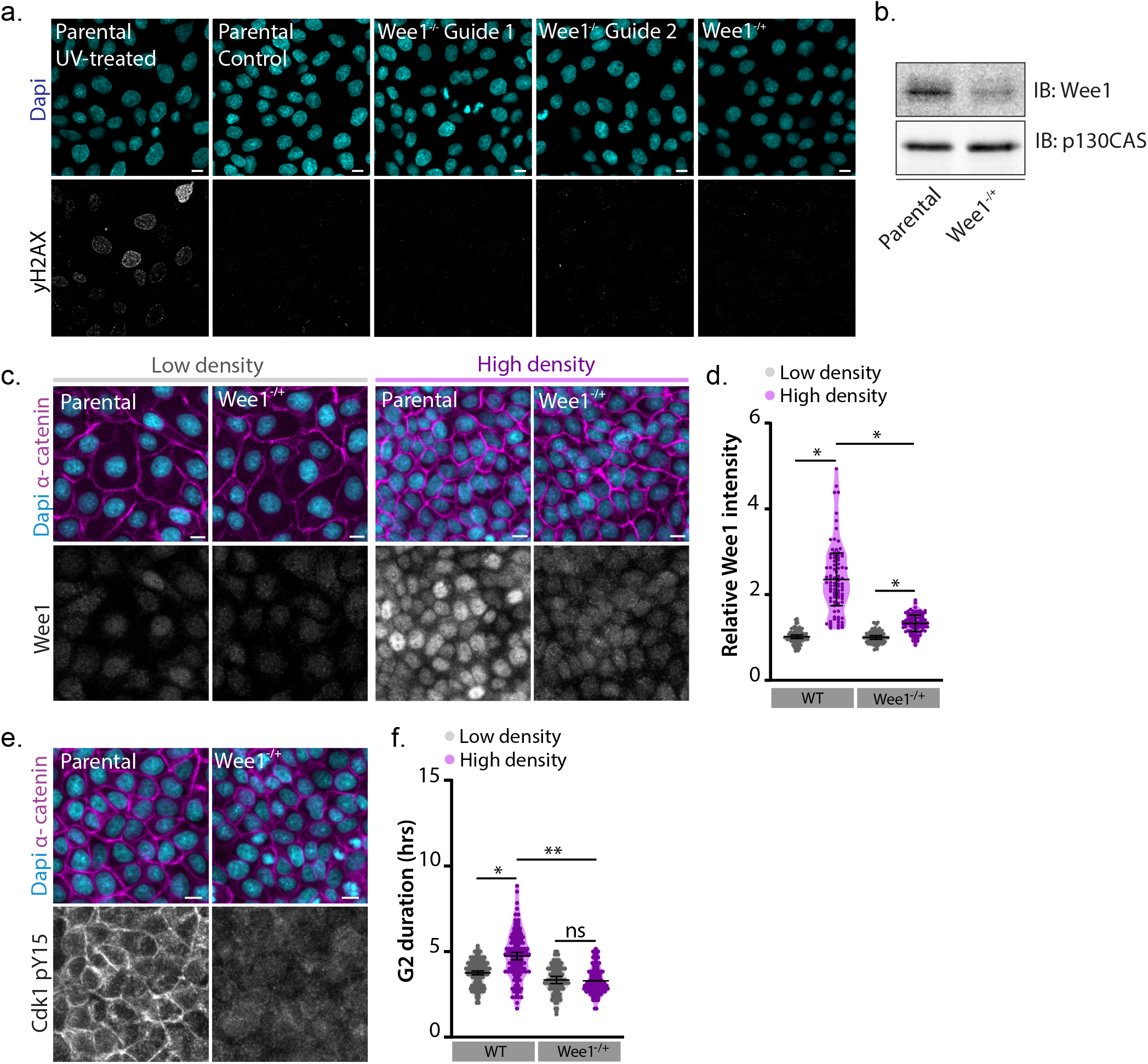
**(a)** Immunostainings of parental MDCK cells (with and without UV irradiation; 20 J/m2), Wee1^−/−^ and Wee1^−/+^ MDCK cells, for the DNA damage marker yH2AX together with Dapi. **(b)** Western blot of lysates from parental and Wee1^−/+^ MDCK cells (grown at high monolayer density) probed for Wee1 and p130CAS. **(c)** Immunostainings of parental and Wee1^−/+^ MDCK cells, grown at low (left) and high (right) monolayer density, for Wee1, together with α-catenin and Dapi. **(d)** Quantification of Wee1 immunofluorescence intensity per cell (normalized to the level of α-catenin intensity) in parental and Wee1^−/+^ MDCK cells grown at low and high monolayer density. n = 90 cells per condition. Data were pooled from 3 independent experiments. Black bars represent the mean and SD of the individual experiments. *P = 0.029 (low density WT vs. high density WT); *P = 0.046 (low density Wee1^−/+^ vs high density Wee1^−/+^); *P = 0.021 (high density WT vs. high density Wee1^−/+^); ratio paired t-test. **(e)** Immunostainings of parental and Wee1^−/+^ MDCK cells for Cdk1 pY15, together with α-catenin and Dapi. **(f)** Quantification of the duration of G2 (hrs), based on expression of mTurquoise-SLBP(18-126), in parental and Wee1^−/+^ MDCK cells grown at low and high monolayer density. n = 120 cells per condition. Data were pooled from 3 independent experiments. Black bars represent the mean and SD of the individual experiments. *P = 0.022; **P = 0.0078, ns = not significant; paired t-test. Scale bars represent 10 μm.

**Supplementary Figure 5.**
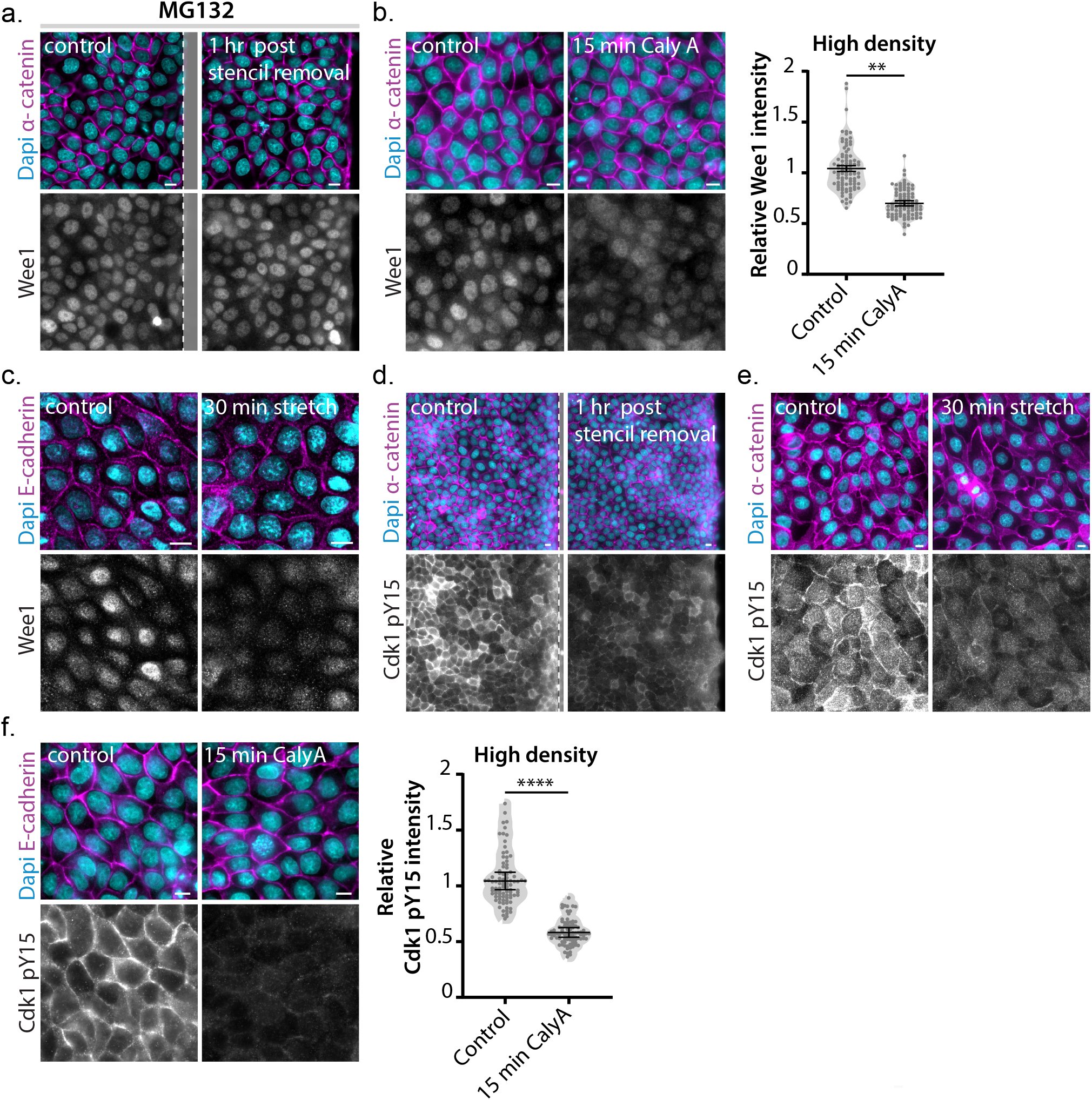
**(a)** Immunostainings of MDCK cells for Wee1, together with α-catenin and Dapi, in an unperturbed high density monolayer and 1 hour after induction of epithelial expansion by removal of the PDMS stencil (from the same experiment shown in Fig. 5a) in presence of the proteasome inhibitor MG132 (5μM). **(b)** Immunostainings of dense MDCK monolayers, treated with the myosin phosphatase inhibitor Calyculin A (15 min; 10 ng/ml) or DMSO control, for Wee1 together with α-catenin and Dapi. The quantification shows Wee1 immunofluorescence intensity per cell (normalized to the level of α-catenin intensity). n = 90 cells per condition. Data were pooled from 3 independent experiments. Black bars represent the mean and SD of the individual experiments. **P = 0.0085; ratio paired t-test. **(c)** Immunostainings of dense control MDCK monolayers, and monolayers subjected to 30 min of 18% uniaxial stretch, for Wee1 together with α-catenin and Dapi. **(d)** Immunostainings of MDCK cells for Cdk1 pY15, together with α-catenin and Dapi, in an unperturbed high density monolayer and 1 hour after induction of epithelial expansion by removal of a PDMS stencil (indicated in grey). **(e)** Immunostainings of dense control MDCK monolayers, and monolayers subjected to 30 min of 18% uniaxial stretch, for Cdk1 pY15 together with α-catenin and Dapi. **(f)** Immunostainings of dense MDCK monolayers, treated with the myosin phosphatase inhibitor Calyculin A (15 min; 10 ng/ml) or DMSO control, for Cdk1 pY15 together with α-catenin and Dapi. The quantification shows Cdk1 pY15 immunofluorescence intensity per cell (normalized to the level of α-catenin intensity). n = 90 cells per condition. Data were pooled from 3 independent experiments. Black bars represent the mean and SD of the individual experiments. ****P = 0.0001; ratio paired t-test. All scale bars represent 10 μm.

**Supplementary Figure 6.**
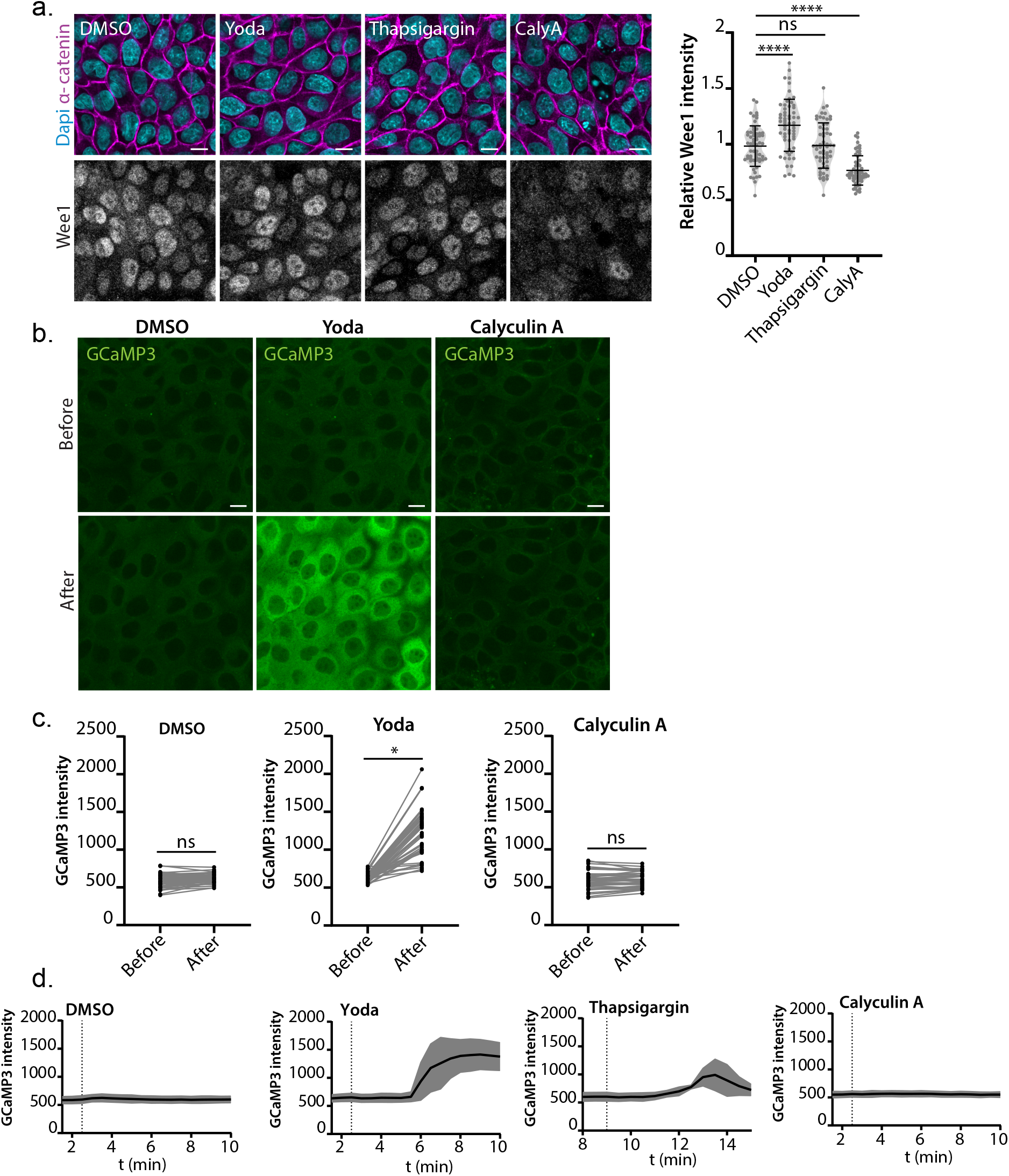
**(a)** Immunostainings of MDCK monolayers treated with DMSO, the Piezo1-selective agonist Yoda (1hr; 10 μM), thapsigargin (Thapsi) (1hr; 1 μM) to induce a calcium influx, or the myosin phosphatase inhibitor Calyculin A (15 min; 10 ng/ml), for Wee1 together with α-catenin and Dapi. The quantification shows Wee1 immunofluorescence intensity per cell (normalized to the level of α-catenin intensity). n = 60 cells per condition. Data were pooled from 2 independent experiments. Black bars represent the mean and SD. ****P <0.0001; ns = not significant; Mann-Whitney. **(b)** Representative still images of MDCK cells stably expressing the GCaMP3 calcium reporter to monitor intracellular calcium levels (Tian et al., 2009), followed over time before and after treatment with DMSO, Yoda (10 μM) and Calyculin A (10 ng/ml). This shows that Calyculin A (which results in Wee1 downregulation, Figs. S5b, S6a) does not induce a calcium influx, in contrast to Piezo1 activation by Yoda. **(c)** Quantification of the GCaMP3 fluorescence signal (mean gray value) per cell in MDCK monolayers before and after treatment with DMSO, Yoda (10 μM) or Calyculin A (10 ng/ml). n = 45 cells per condition. Data were pooled from 3 independent experiments. *P = 0.026; ns = not significant; paired t-test. **(d)** Representative traces of the fluorescence intensity (mean gray value) of MDCK GCaMP3 cells, followed over time before and after treatment with DMSO, Yoda (10 μM), thapsigargin (1 μM) or Calyculin A (10 ng/ml). The black line indicates the mean fluorescence signal, with the SD indicated in grey. Scale bars represent 10 μm.

